# Quantifying Tensile Forces at Cell–Cell Junctions with a DNA-based Fluorescent Probe

**DOI:** 10.1101/2020.01.07.897249

**Authors:** Bin Zhao, Ningwei Li, Tianfa Xie, Chungwen Liang, Yousef Bagheri, Yubing Sun, Mingxu You

## Abstract

Cells are physically contacting with each other. Direct and precise quantification of forces at cell–cell junctions is still challenging. Herein, we have developed a DNA-based ratiometric fluorescent probe, termed DNAMeter, to quantify intercellular tensile forces. These lipid-modified DNAMeters can spontaneously anchor onto live cell membranes. The DNAMeter consists of two self-assembled DNA hairpins of different force tolerance. Once the intercellular tension exceeds the force tolerance to unfold a DNA hairpin, a specific fluorescence signal will be activated, which enables the real-time imaging and quantification of tensile forces. Using E-cadherin-modified DNAMeter as an example, we have demonstrated an approach to quantify, at the molecular level, the magnitude and distribution of E-cadherin tension among epithelial cells. Compatible with readily accessible fluorescence microscopes, these easy-to-use DNA tension probes can be broadly used to quantify mechanotransduction in collective cell behaviors.

## INTRODUCTION

Intercellular mechanical forces, especially tensile forces, play important roles in development, tissue healing and cancer invasion^1–3^. These tensile forces at cell–cell junctions actively reshape the tissues during morphogenesis in embryos, and are also evident in quiescent adult tissues^4–6^, especially epithelial and endothelial monolayers^7,8^. Cadherins constitute a superfamily of cell–cell adhesion molecules that are expressed in various types of cells^9,10^. It is known that cadherins can sense and mediate tensile forces at cell–cell junctions^11^, which are required for several cellular functions and the organizations of soft tissues^12–14^. These intercellular cadherin-mediated forces are shown to regulate cellular homeostasis and collective migration during embryo development, wound healing, and pulmonary system homeostasis^6,15–17^. Elucidating the mechanisms of force sensing and transduction in cadherins is therefore critical for revealing the fundamental principles in the collective organizations and motions of a cell population^18,19^. While single cell studies clearly revealed the force-sensing capability of cadherin molecules, mapping the spatiotemporal dynamics of their forces in situ require precise, real-time measurement of tension at cell–cell junctions^20–22^.

Generally, two strategies are currently available to estimate intercellular forces. Monolayer stress microscopy utilizes cell–matrix traction force data to deduce mechanical forces at cell–cell junctions, with the assumption that total forces experienced by each cell remain zero^23^. Traction force microscopy has been used to elucidate the relationships between the total cellular forces on extracellular matrix and the endogenous intercellular forces^54^. However, these methods can only be applied to a monolayer of cells. It requires extensive image analysis and data processing. Moreover, the force deduced is not specific for certain junctional molecules. Similarly, intercellular forces between a pair of cells have been measured by microfabricated cantilever pillars^13^. However, in addition to the above-mentioned drawbacks, these cantilever pillars can only measure forces between a pair of cells, one at a time, and require advanced microfabrication facilities.

In another strategy, genetically encoded protein-based tension probes have been developed to measure intercellular forces mediated by cadherins or platelet endothelial cell adhesion molecule^8,24–26^. However, the routine use of these sensors is still limited due to their labor-intensive design and validation. The functions of many junctional proteins will be disrupted after insertion of a large protein sensor (~500 amino acids). The small force measurement range (1–12 pN) and low sensitivity of fluorescence signals of these probes (~10-fold lower than sensors using common organic dyes)^27^ further hinders the widespread applications of these genetically encoded probes.

We have recently developed a DNA-based probe to visualize intercellular tensile forces at cell–cell junctions^28^. In this system, a pair of cholesterol anchors was used to insert this DNA hairpin-based probe onto live cell membranes^29,30^. Once the intercellular tensile force exceeds the threshold value to unfold the DNA hairpin, the separation of a fluorophore-quencher pair results in the activation of fluorescence signals. These DNA probes are well suited for intercellular force measurement. First, the probes function simply by incubation with target cells. There is no need for cloning or transfection. Secondly, different mechanosensitive ligands or receptors can be directly conjugated within these DNA probes, which allows the facile study of specific junctional molecule-mediated force transduction. Thirdly, by tuning the sequence and duplex length of the DNA hairpin, the force tolerance of the probe can be rationally adjusted in a large range^31–33^. Moreover, a broad choice of organic fluorophores and quenchers allows highly sensitive imaging of tensile forces.

However, these DNA tension probes still have several limitations. For example, to quantify the intercellular forces, the heterogeneous membrane distribution of the probes should be first normalized. Moreover, each DNA hairpin unfolds in a narrow threshold force range (± 2 pN)^34^, multiple probes of different threshold values are needed to measure a broad range of intercellular forces. In addition, many collective cell behavior studies require a long-term measurement of intercellular forces^17,35–37^. However, these DNA probes have a limited anchoring persistence on the cell membranes (~ 2–4 h). To overcome these limitations, in this study we have developed a second-generation <u>DNA</u>-based <u>me</u>mbrane <u>te</u>nsion ratiometric probe, termed DNAMeter.

The DNAMeter was designed to be highly adaptable, consisting of two self-assembled DNA hairpins with different threshold forces and a lipid tail to anchor onto live cell membranes (Fig. 1). To quantify the intercellular tension based on the fluorescence signal, an internal reference fluorophore was introduced to normalize the membrane distribution of the DNAMeter. In addition, two orthogonal fluorophore-quencher pairs were conjugated at the end of each DNA hairpin to report different magnitudes of forces. By measuring each reporter-to-reference fluorescence intensity ratio, molecular scale intercellular force distributions can be quantified at cell–cell junctions. Using E-cadherin-mediated intercellular tension as an example, we demonstrated here a simple and general approach to quantify mechanical characteristics of collective cell behaviors.

**Fig. 1.**
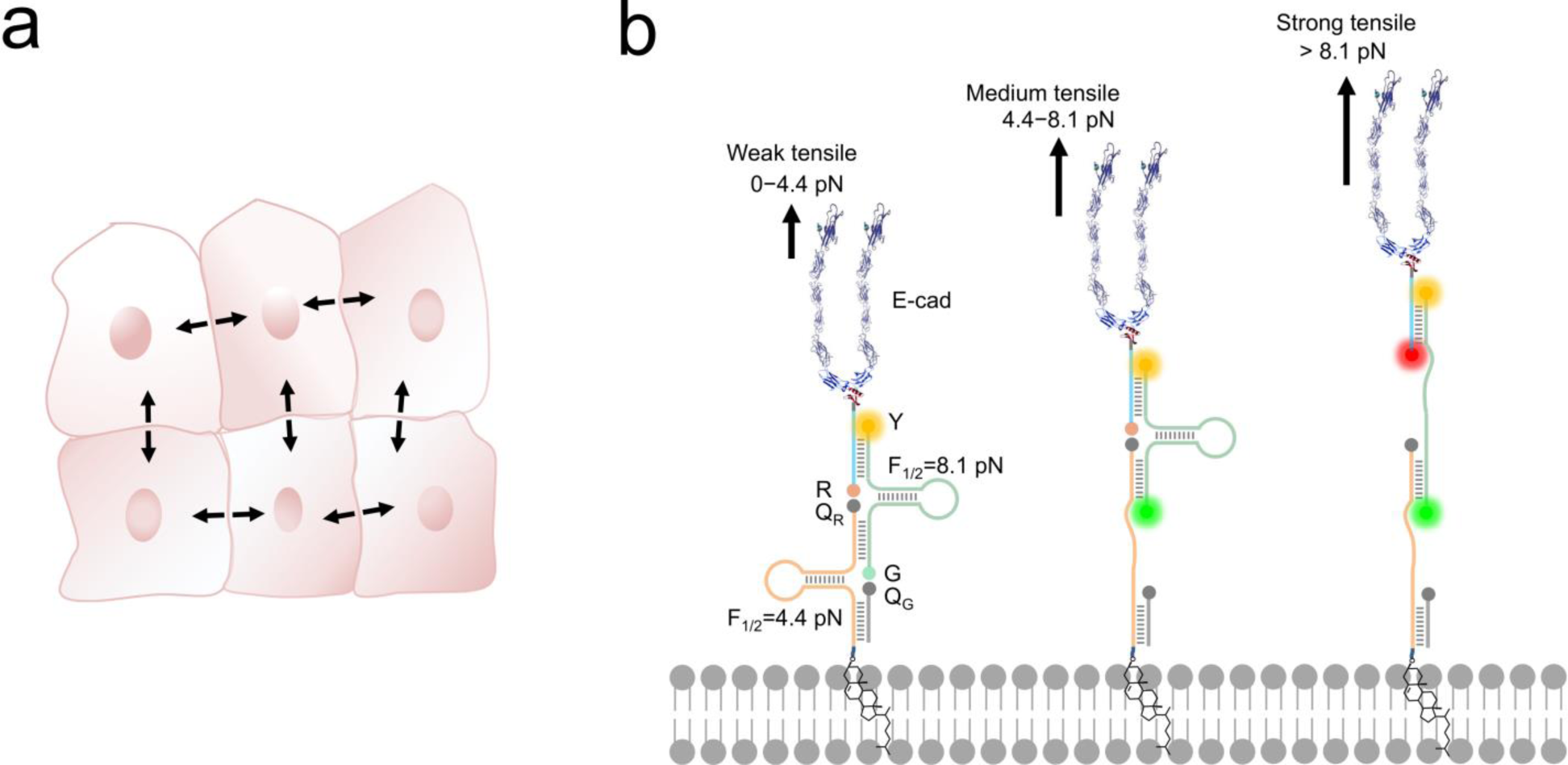
Design of the DNAMeter to quantify tensile forces at cell–cell junctions. (**a**) Schematic of collective cell system experiencing intercellular tensile forces. Black arrows indicate the forces at cell–cell junctions that we are studying in this project. (**b**) The construction of an EC-DNAMeter on a live cell membrane. The DNAMeter is comprised of a cholesterol-modified 22%GC DNA hairpin strand (orange, F_1/2_= 4.4 pN), a 66%GC hairpin strand (light green, F_1/2_= 8.1 pN), a ligand strand (light blue) and a helper strand (grey). The DNA strands was further modified with E-cadherin (E-cad) through a Protein G linker to form the EC-DNAMeter. Upon experiencing different magnitudes of tensile forces as generated by the neighboring cells, the FAM (G) and/or Cy5 (R) fluorophore separates from the corresponding quencher, Dabcyl (Q_G_) and/or QSY^®^21 (Q_R_). Here, a TAMRA fluorophore (Y) acts as the internal reference for the ratiometric imaging and quantification.

## RESULTS AND DISCUSSION

### Design and characterization of the DNAMeter

The DNAMeter is designed based on the self-assembly of four oligonucleotide strands (Fig. 1b and Table S1). Two of the strands contain a 25-nucleotide-long DNA hairpin with 22% and 66% G/C base pairs to detect weak and strong tensile force, respectively. As an internal reference, a TAMRA dye (λ_ex_/λ_em_: 557/579 nm, denoted as Y) was modified at one end of the 66%GC DNA hairpin strand. To detect the folding/unfolding switch of the 66%GC DNA hairpin, a Cy5-QSY^®^21 fluorophore-quencher pair (λ_ex_/λ_em_: 640/659 nm, denoted as R-Q_R_) was conjugated next to the end of this hairpin. Similarly, a FAM-Dabcyl fluorophore-quencher pair (λ_ex_/λ_em_: 488/519 nm, denoted as G-Q_G_) was used to measure the folding/unfolding of the 22%GC DNA hairpin. After calculating the unfolding free energy of hairpin at zero force and the free energy of stretching the corresponding single stranded DNA^32,34^, based on a worm-like chain mode^38,39^, we have determined the tensile force threshold (F_1/2_) of the 22%GC and 66%GC DNA hairpin to be 4.4 pN and 8.1 pN, respectively (Methods and Table S2). Here, F_1/2_ is defined as the force at which the DNA hairpin has 50% probability of being unfolded.

When experiencing a weak tensile force (<4.4 pN), both FAM-Dabcyl and Cy5-QSY^®^21 pairs remain at close proximity, resulting in low fluorescence level in both reporter channels (denoted as G-/R-). In contrast, a strong tensile force (>8.1 pN) results in the stretching out of both 22%GC and 66%GC hairpins. Both fluorophores will separate from the corresponding quencher, leading to an increase in both FAM and Cy5 fluorescence signal (denoted as G+/R+). In another case, a medium tensile force (4.4–8.1 pN) opens up the 22%GC hairpin, but not the 66%GC hairpin, so only the FAM signal will be activated (denoted as G+/R-). As a result, we can image different ranges of tensile forces based on the two reporter channels.

To test the efficiency of this probe design, we prepared an E-cadherin-modified DNAMeter (termed EC-DNAMeter) to quantify E-cadherin-mediated intercellular tensile forces at the Madin-Darby canine kidney (MDCK) epithelial cell–cell junctions. The EC-DNAMeter was prepared using a Protein G linker to couple the IgG/Fc-fused E-cadherin with the DNAMeter (Methods). Compared with direct chemical conjugation, the Protein G linker helps to avoid the loss of E-cadherin activities^28,40^. In addition, to allow the probe to insert onto MDCK cell membranes, a cholesterol anchor was conjugated at the other end of the DNAMeter.

After demonstrating the formation of the DNAMeter in a gel mobility shift assay (Fig. S1), the cell membrane insertion efficiency of the DNAMeter was studied. Here, we prepared a non-quenched DNAMeter (nqDNAMeter) by using DNA strands that are not modified with Dabcyl or QSY^®^21 quencher. The fluorescence of the nqDNAMeter is always on and is independent of intercellular forces. As a result, the cell membrane fluorescence intensity can be used to indicate the concentration of the immobilized probes. Indeed, obvious fluorescence signal on MDCK cell membranes was shown shortly after adding these nqDNAMeter probes (Fig. S2).

We have further studied the membrane anchoring efficiency of the DNAMeter containing one or two cholesterol tail. Interestingly, one cholesterol-modified nqDNAMeter (1Chol-nqDNAMeter) exhibited higher insertion efficiency (2.1-fold) on MDCK cell membranes than the more hydrophobic two cholesterol-modified one (2Chol-nqDNAMeter) (Fig. S2). This might be due to the relatively larger critical micelle concentration value of the 1Chol-nqDNAMeter as compared to 2Chol-nqDNAMeter^41^. As a result, more monomeric nqDNAMeter could exist in the solution when one cholesterol was anchored. Indeed, our recent data indicated that the cell membrane anchoring of the lipid-DNA conjugates stems mainly from the monomeric form, instead of the aggregation form^55^. Previous studies have suggested that ~100 pN tensile force is required to extract a cholesterol from lipid bilayers^42^. As a result, the membrane insertion of the cholesterol should be quite stable during the unfolding of DNA hairpins (4.4 pN and 8.1 pN). Unless specifically indicated, one cholesterol-based construct was used for the following studies.

One potential concern of the DNAMeter probe is that it may be activated by the *cis* receptor–ligand interactions on the same cell membrane, rather than between neighboring cells. To study if these DNA probes prefer to “stand” (favoring *trans* interactions) or “lie down” (favoring *cis* interactions) on membrane surfaces, we performed atomistic molecular dynamics simulation in a DNAMeter/lipid bilayer membrane system (Supplementary Note 1). Our simulation results indicated that the tilting angle (θ) of the DNAMeter with respect to the membrane surface is always within 30º (see Fig. S3). Indeed, these membrane-anchored DNA probes prefer to “stand” (θ< 30º) on the membrane surface and favor the sensing of *trans* interactions between cells. The reason for DNA probes to maintain such orientation is likely attributed to the electrostatic repulsion between DNA strands and cell membranes, which are both negatively charged.

We next asked if we could distinguish the unfolding and the folding state of DNA hairpins based on their fluorescence intensities. To determine the fluorescence of the unfolded DNAMeter, we prepared a de-quenched probe (dqDNAMeter) by incubating the DNAMeter with DNA strands that are complementary to the 22%GC and 66%GC hairpins, respectively (Fig. S4). Based on the fluorescence intensity ratio between the DNAMeter and dqDNAMeter, in the absence of external forces, the quenching efficiency for Cy5 and FAM in the DNAMeter was measured to be 81% and 70%, respectively (Fig. 2a and 2b). Meanwhile, the DNAMeter and dqDNAMeter exhibited the same TAMRA fluorescence intensity, which can act as a standard reference to normalize probe concentrations (Fig. 2c). In addition, after incubating 1 µM DNAMeter or dqDNAMeter with MDCK cells for 30 min, similarly, almost the same TAMRA fluorescence was observed. In contrast, 6.7-fold and 3.1-fold activation of Cy5 and FAM fluorescence exhibited after the unfolding of DNA hairpins (Fig. S5). These results indicate that the folding and unfolding of 22%GC and 66%GC hairpins indeed can be visualized based on changes in the fluorescence intensity.

**Fig. 2.**
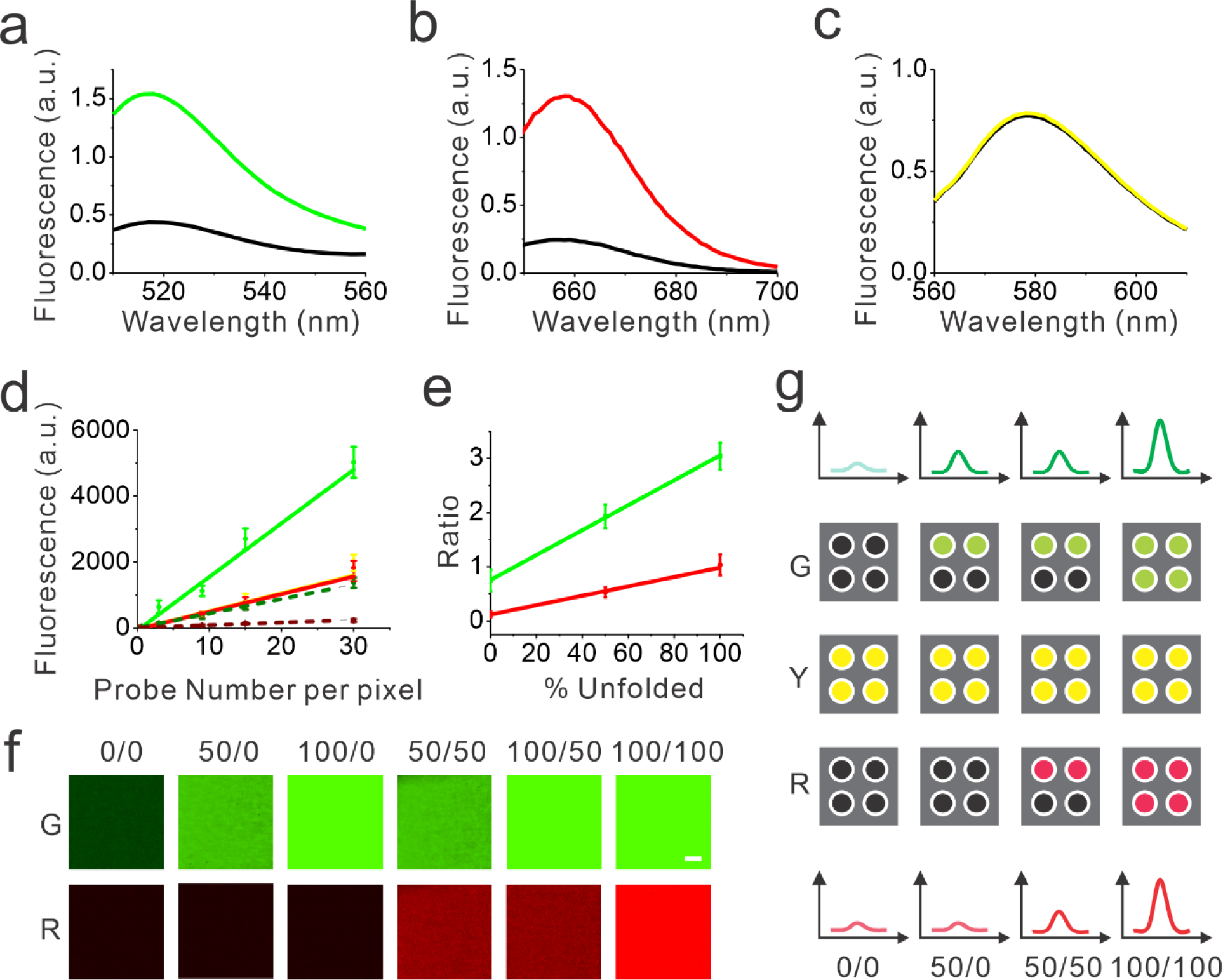
In vitro characterization of the DNAMeter. (**a – c**) The fluorescence spectra of the EC-DNAMeter (color line) and de-quenched EC-DNAMeter (black line) was measured in terms of FAM (a), Cy5 (b), and TAMRA (c). The excitation wavelength was 488 nm, 630 nm, and 550 nm, respectively. (**d**) Calibration curves to correlate the membrane fluorescence intensity with the number of probes per pixel on a supported lipid monolayer. The de-quenched DNAMeter was used to measure the calibration curves for unfolded 22%GC hairpin (green solid line), 66%GC hairpin (red solid line), and TAMRA reference (yellow solid line). While the DNAMeter was used to calibrate for the folded 22%GC hairpin (dark green dashed line) and 66%GC hairpin (dark red dashed line). (**e**) Correlation of the G/Y or R/Y ratio with the percentage of unfolded hairpins in individual pixels. G/Y (green line) indicates the percentage of unfolded 22%GC hairpins. R/Y (red line) indicates the percentage of unfolded 66%GC hairpins. (**f**) Fluorescence images by adding different combinations of the DNAMeter and de-quenched DNAMeter onto a supported lipid monolayer. For example, 100/50 means that 22%GC and 66%GC DNA hairpins were 100% and 50% unfolded, respectively. Scale bar, 5 µm. (**g**) Schematic of the correlation between the fluorescence intensity and the number of unfolded DNA hairpins in each pixel. The top and bottom spectra illustrate the fluorescence intensity of FAM and Cy5. Each square indicates an individual pixel, and each dot represents a single DNA probe. For example, 50/0 indicates that the percentage of unfolded 22%GC and 66%GC DNA hairpins is 50% and 0%, respectively.

### Calibration of the DNAMeter on supported lipid monolayers

We next asked if we could further quantify the percentage of unfolded DNA hairpins based on the fluorescence intensities. For this purpose, we prepared a supported lipid monolayer system using soybean polar extract^43^. Cholesterol-modified DNAMeter can anchor into this monolayer and diffuse freely^29^. By mixing the soybean polar extract with different amount of DNA probes, we can precisely control the membrane density of the DNAMeter on lipid monolayers. After preparing a series of monolayers with different probe densities, we measured the corresponding membrane fluorescence intensity with a spinning disk confocal fluorescence microscope. The same setup and parameters of the microscope was used for the following cellular measurements as well.

The obtained fluorescence intensities were then plotted as a function of probe densities for the calibration. A linear correlation between the fluorescence intensity and the DNAMeter concentration was observed for all the fluorophores, including FAM (G), Cy5 (R), and TAMRA (Y) (Fig. 2d and 2e). Similarly, a linear correlation was observed with all these fluorophores in the dqDNAMeter as well (Fig. 2d and 2e). After subtracting the background fluorescence for each channel, the fluorescence intensity ratio of both FAM/TAMRA (G/Y) and Cy5/TAMRA (R/Y) is independent on the probe concentration due to the linear correlation between the probe density and fluorescence (Fig. 2d). Such concentration-independent G/Y and R/Y ratio was observed with both the DNAMeter and dqDNAMeter, while the dqDNAMeter exhibited a 4.0-fold and 8.3-fold higher intensity ratio. The G/Y and R/Y ratio can thus be used to quantify the membrane dqDNAMeter-to-DNAMeter probe density ratio, as well as the percentage of unfolded 22%GC and 66%GC DNA hairpins, respectively.

Our next goal is to validate if the G/Y and R/Y ratio can be used to quantify the percentage of unfolded DNA hairpins. We prepared mixtures of dqDNAMeter and DNAMeter, with a ratio of 0:1, 0.5:0.5, and 1:0. Indeed, both G/Y and R/Y ratio are linearly correlated with the percentage of the unfolded dqDNAMeter (Fig. 2e). We have also tested if the G/Y and R/Y ratio can orthogonally report the unfolding of 22%GC and 66%GC DNA hairpins, respectively. By adding only a complementary DNA strand to either 22%GC or 66%GC DNA hairpin, we prepared de-quenched DNAMeter with only one hairpin being unfolded. After mixing different ratios of these two partially unfolded DNAMeter, indeed, the FAM and Cy5 signal can be used to quantify the amount of unfolded 22%GC or 66%GC DNA hairpin, without interfering with each other (Fig. 2f). All these results indicated that we could quantify the unfolding of DNA hairpins in the DNAMeter by measuring the G/Y and R/Y ratio. Based on the standard calibration curve (Fig. 2d), we can also use the TAMRA fluorescence to quantify the number of probes per individual pixel of images. As a result, we can quantitatively determine not only the percentage, but also the number of unfolded DNA hairpins from the images (Fig. 2g).

### Imaging and quantification of E-cadherin-mediated tensile forces

Before imaging intercellular forces, we wondered if the addition of DNAMeter would impair the adhesion and mechanical function of cell–cell junctions. We first studied the effect of membrane-anchored EC-DNAMeter on the force-dependent recruitment of vinculin to the adherens junctions^56,57^. Immunofluorescence staining was used to image the cellular locations of vinculin in MDCK cells before and after adding the DNAMeter. MDCK cells have been widely used as a model cell line to study E-cadherin-mediated tensile forces^44^. No significant difference in the junction vinculin fluorescence was observed (Fig. S6). We have also used western blot to study the effect of DNAMeter on the membrane expression of another critical cell–cell adhesion protein, β-catenin^58,59^. Again, the amount of β-catenin in MDCK cell membranes was quite similar in the presence or absence of EC-DNAMeter anchoring (Fig. S6). As a result, the addition of DNAMeter will not influence the mechanotransduction at cell–cell junctions.

We next applied the EC-DNAMeter to image E-cadherin-mediated intercellular tensile forces at MDCK cell–cell junctions. After incubating the pre-assembled EC-DNAMeter with MDCK cells for 1 h, the cell membrane fluorescence signal of FAM (λ_ex_/λ_em_: 488/530 nm), TAMRA (λ_ex_/λ_em_: 561/590 nm), and Cy5 (λ_ex_/λ_em_: 640/675 nm) were imaged with a spinning disk confocal microscope (Fig. 3a). Here, we denoted the fluorescence of FAM, TAMRA, and Cy5 as G, Y, and R, respectively. For a given DNAMeter-modified cell membrane (Y+), the weak (<4.4 pN), medium (4.4–8.1pN), and strong (>8.1 pN) E-cadherin-mediated intercellular tensile forces can be visualized based on the fluorescence distribution of G-/R-, G+/R-, and G+/R+, respectively. A large number of G+/R- and some G+/R+ pixels were clearly observed at MDCK cell–cell junctions (Fig. 3a). To test if the fluorescence activation is indeed mediated by E-cadherin interactions, we prepared a control DNAMeter without the modification of E-cadherin, denoted as ProG-DNAMeter. As expected, limited FAM and Cy5 fluorescence was observed, while the TAMRA fluorescence was similar as that of the EC-DNAMeter. These results indicated that we could visualize E-cadherin-mediated tensile forces using the EC-DNAMeter.

**Fig. 3.**
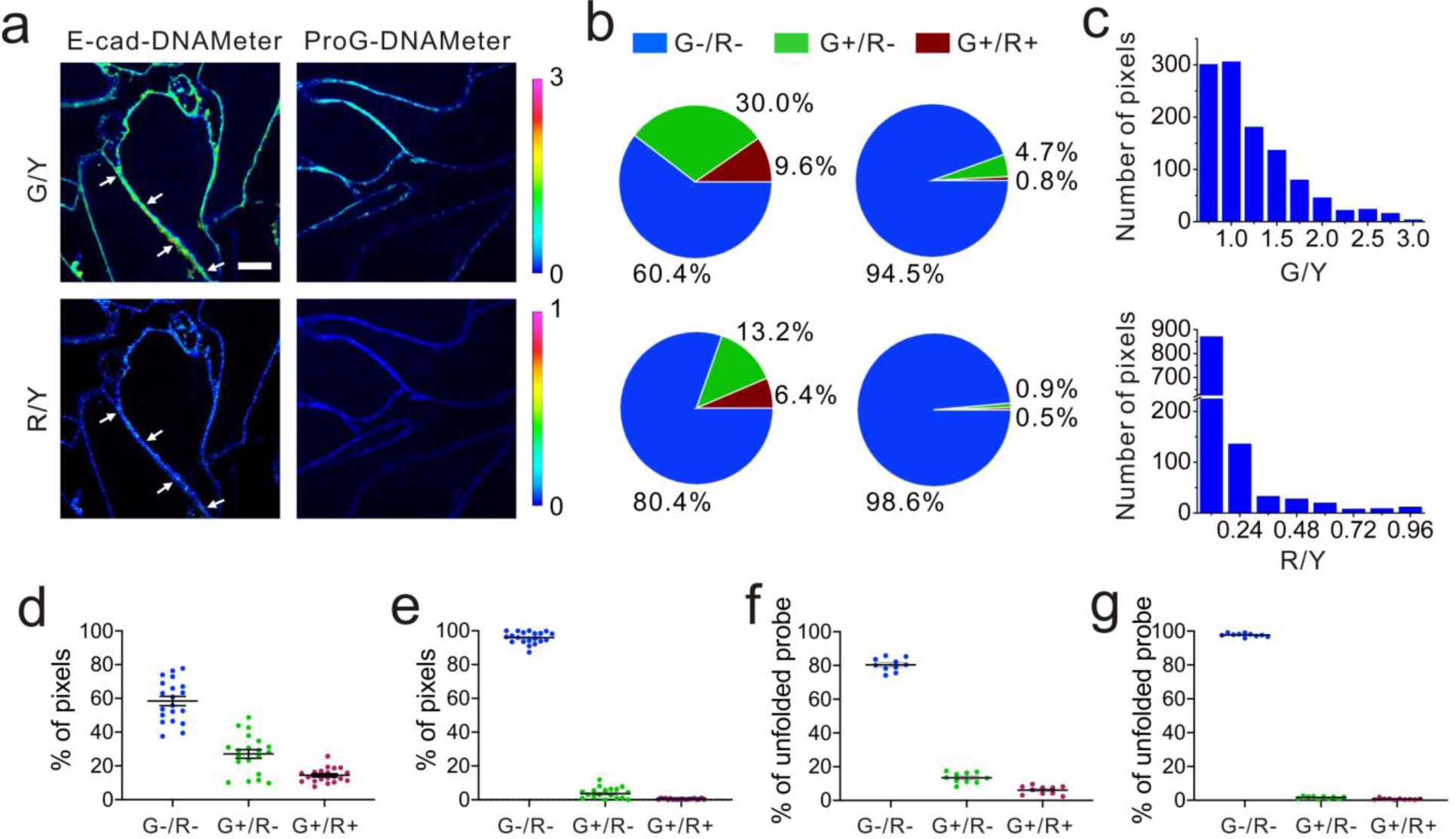
Quantification of E-cadherin-mediated tensile forces at MDCK cell–cell junctions. (**a**) Representative fluorescence images of MDCK cells after inserting the EC-DNAMeter. G/Y stands for the fluorescence ratio of FAM to TAMRA, indicating the tensile forces above 4.4 pN. R/Y is the fluorescence ratio of Cy5 to TAMRA, indicating the tensile forces above 8.1 pN. The ProG-DNAMeter that lacks E-cad modification is used as a control. Scale bar, 5 µm. (**b**) Quantitative analysis of the tension based on the fluorescence images. The top panels show the percentage of pixels experiencing forces as quantified with the EC-DNAMeter (left) and ProG-DNAMeter (right). The bottom panels indicate the percentage of unfolded probes with the EC-DNAMeter (left) and ProG-DNAMeter (right). G+ (or G-) indicates the fluorescence ratio of FAM to TAMRA is above (or below) the threshold, respectively. Similarly, R+ (or R-) indicates the fluorescence ratio of Cy5 to TAMRA is above (or below) the threshold value. (**c**) The distribution of pixels within different subranges of G/Y or R/Y ratios for the representative junction denoted by white arrows in the panel (a). (**d, e**) Statistical analysis of tensile force distributions in terms of the percentage of pixels at different cell–cell junctions (N= 20) with the (d) EC-DNAMeter or (e) ProG-DNAMeter. (**f, g**) Statistical analysis of tensile force distributions in terms of the percentage of unfolded probes at different cell–cell junctions (N= 10) with the (f) EC-DNAMeter or (g) ProG-DNAMeter.

We next asked if we could quantify the distribution of different magnitudes of tensile forces at cell–cell junctions. Here, we quantified the force distribution by either the number of bright pixels or the number of unfolded DNA probes. To calculate the number or percentage of pixels considering as G+ or R+, we first measured the membrane statistical fluorescence distribution of the negative control, the ProG-DNAMeter (Fig. 2e). A threshold value of G/Y> 1.0 and R/Y> 0.24 was determined to distinguish the pixels experiencing tensile forces above 4.4 pN and 8.1 pN, respectively. After counting the total number of probe-immobilized pixels (Y+) at cell–cell junctions, we quantified the percentage of junction pixels experiencing the tensile forces (Fig. 3b). For example, at a representative cell–cell junction (Fig. 3a), the weak (<4.4 pN), medium (4.4–8.1pN), and strong (>8.1 pN) tension was present in ~60.4%, 30.0%, and 9.6% membrane areas, respectively. We have further calculated these distributions at another 20 cell–cell junctions. On average, under the studied condition when MDCK cells were stably adhered to each other, the fraction of pixels experiencing the weak, medium, and strong tension was 58.5±12.3%, 27.1±11.5% and 14.4±4.2%, respectively (Fig. 3d and Table S3).

The second approach to quantify the force distributions is based on the number and percentage of the unfolded EC-DNAMeter. As mentioned above, the percentage of unfolded 22%GC and 66%GC DNA hairpin in each pixel can be calculated by measuring the G/Y and R/Y ratio (Fig. 2g). The number of unfolded probes can then be quantified based on the TAMRA fluorescence and the standard calibration curve (Fig. 2f). Our data indicated that 13.2±3.1% and 6.4±2.4% EC-DNAMeter probe was unfolded by 4.4–8.1 pN and >8.1 pN tension, respectively, while 80.4±3.9% probe remained folded (Methods, Fig. 3b and 3c). As a control, unfolding of the ProG-DNAMeter was negligible (Fig. 3g).

When comparing the obtained data from two approaches, we found that by measuring the percentage of unfolded DNA probes, force distributions at different cell–cell junctions were more homogeneous (Fig. 3d and 3f). Considering these MDCK cells were experiencing similar physical environment and cell–cell adhesions, it may be more accurate to determine force distributions by measuring the percentage of unfolded probes rather than that of bright pixels. Even though the percentage of bright pixels is easier to be quantified, the accuracy of this approach is influenced by the choice of threshold values, e.g., G/Y> 1.0 and R/Y> 0.24 in this case. Meanwhile, even in pixels that are brighter than the threshold value, many DNAMeters can be still in the folded form. In contrast, the percentage of unfolded DNA hairpins can more accurately report, at the molecular level, the force distributions experienced by the DNAMeter.

### Dynamics of E-cadherin-mediated tension

To validate if the EC-DNAMeter indeed measured E-cadherin-mediated tensile forces, we have further studied the effect of ethylene glycol-bis (β-aminoethyl ether)-N,N,N′,N′-tetraacetic acid (EGTA) treatment on MDCK cell adhesions. E-cadherin interactions are gated by extracellular Ca^2+^ ions that can rigidify the extracellular domains of cadherins and promote cadherin–cadherin junctional interactions^45^. As a selective Ca^2+^ chelating agent, EGTA can disrupt E-cadherin interactions at cell–cell junctions^46^. Indeed, after the insertion of the EC-DNAMeter onto MDCK cell membranes, the addition of EGTA triggered a rapid and substantial loss of the fluorescence signal, accompanied with cell dissociations (Fig. 4a and S7).

**Fig. 4.**
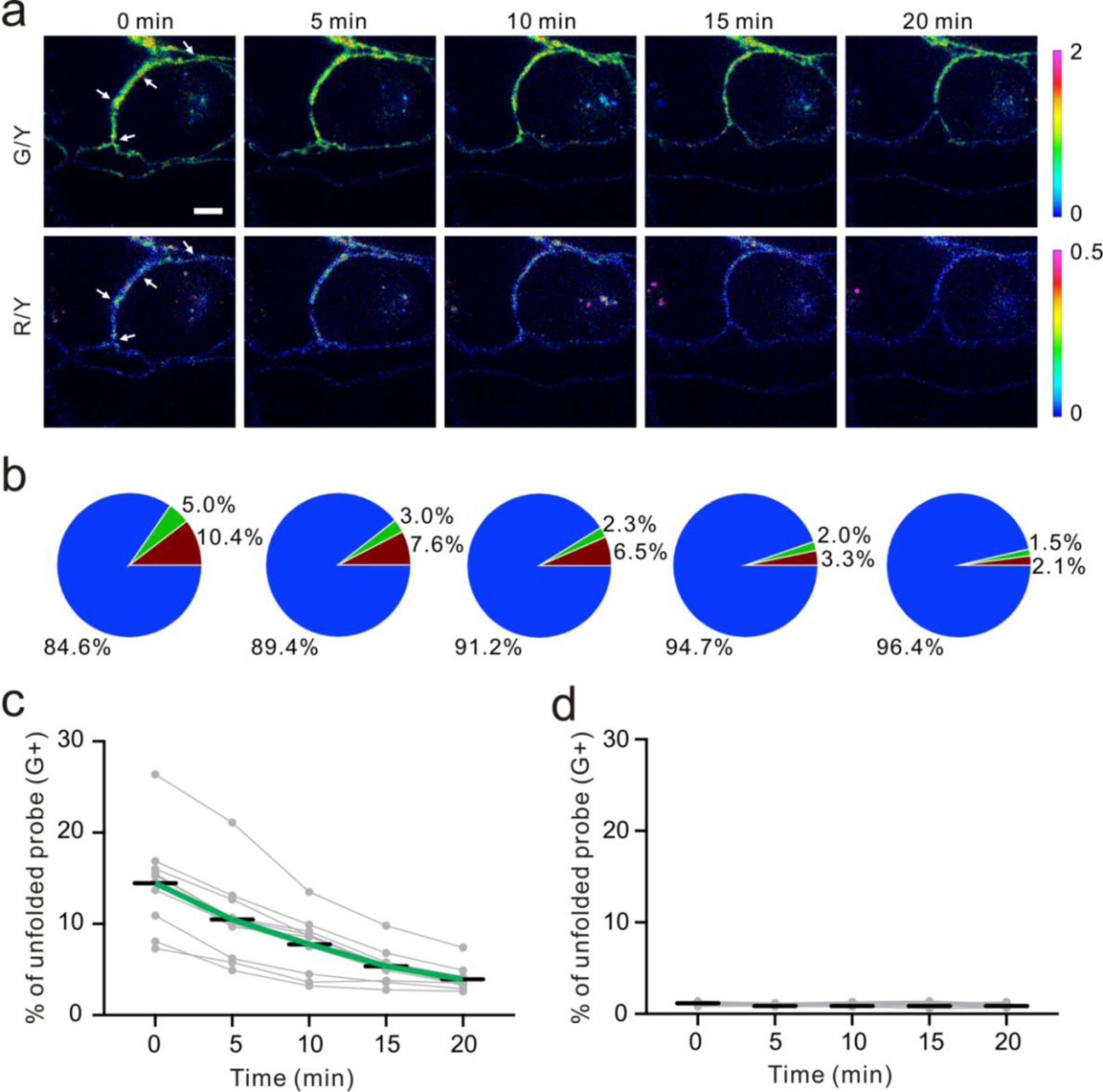
Dynamics and disruptions of the E-cadherin-mediated tension. (**a**) Fluorescence images of EC-DNAMeter-modified MDCK cells after adding 10 mM EGTA at 0 min. The cell–cell junction denoted by white arrows was used for the quantitative analysis in the panel (b). Scale bar, 5 µm. (**b**) The quantitative analysis of tension revealed by percentage of unfolded probes after adding EGTA. Each pie chart corresponds to the images above it in the panel (a). The blue, green, and red region indicted the distribution of tensile forces in the range of <4.4 pN, 4.4–8.1 pN, and >8.1 pN, respectively. (**c, d**) Statistical analysis of the dynamic changes in the intercellular tensile forces (>4.4 pN) at different cell–cell junctions (N= 10) with the (c) EC-DNAMeter or (d) ProG-DNAMeter.

We next asked if the EC-DNAMeter could be used to monitor the dynamic variations of E-cadherin-mediated intercellular tension after the EGTA treatment. Indeed, at a representative cell–cell junction, within 20 min after adding 10 mM EGTA, the percentage of medium tension (4.4–8.1 pN) gradually decreased from 5.0% to 1.5%, and meanwhile large tension (>8.1 pN) dropped from 10.4% to 2.1% (Fig. 4a and 4b). Further quantification of more cell–cell junctions confirmed that the EC-DNAMeter could be used to measure the dynamics of intercellular E-cadherin tension. Interestingly, a linear decrease in the number of membrane probes experiencing medium or large tensile forces (>4.4 pN) was observed after adding EGTA, with a rate constant ~53 µm^−2^ min^−1^ (Fig. 4c). In a control experiment, a constant unfolding percentage of the ProG-DNAMeter (~1.1%) was shown before and after adding EGTA (Fig. 4d, S8, and S9).

As another example, we applied the EC-DNAMeter to monitor the ML-7-induced changes in the E-cadherin tension. ML-7 can inhibit the activity of myosin light chain kinase and impair the ability of cells to concentrate E-cadherin at cell–cell junctions^60^. As expected, the treatment of ML-7 induced a gradual decrease in the G/Y and R/Y ratio at MDCK cell–cell junctions (Fig. S10). For example, at a representative junction, after adding 100 µM ML-7, the percentage of large tension (>8.1 pN) gradually decreased within 20 min from 6.3% to 0.5%, and meanwhile medium tension (4.4–8.1 pN) dropped from 10.2% to 0.5% (Fig. S10). The statistical analysis of more cell–cell junctions further confirmed this observation (Fig. S11). In contrast, the control probe, ProG-DNAMeter, displayed a constant unfolding percentage at ~0.8% (Fig. S10 and S11). Indeed, the EC-DNAMeter can be used to study the dynamic E-cadherin tensions at cell–cell junctions.

### Force mapping during collective cell migration

Cooperative intercellular forces drive cellular motions and play vital roles in collective cell migration^23,47^. We asked if the DNAMeter could be used to quantify intercellular tensions during collective migration of an epithelial monolayer. Epithelial migration occurs on a time scale of hours-to-days. We first wondered if the DNAMeter allows long-term force measurement. For the sake of simplicity, we prepared a non-quenched EC-DNAMeter containing only a 22%GC DNA hairpin (nqEC22-DNAMeter). After incubating this nqEC22-DNAMeter with a confluent MDCK cell monolayer for 1 h, the probe persistence on the cell membrane was studied. Our results indicated that in a complete growth medium, the cell membrane fluorescence would completely disappear within 3 h (Fig. S12). Since the growth medium is needed for collective epithelial migrations, the DNAMeter cannot be directly used for the force mapping.

To achieve a long-term force measurement, we asked if the cell membrane probe density could be recovered by simply adding fresh DNAMeter. To test that, we first anchored 0.2 µM nqEC22-DNAMeter onto MDCK cell membrane in HEPES-buffered saline, followed by replacing with complete growth medium. After 3 h incubation, almost no fluorescence was observed on the cell membrane. By further replacing the growth medium with buffer containing 0.2 µM of fresh nqEC22-DNAMeter, again, strong fluorescence and a similar level of membrane probe density (95.6±4.5%) was observed at cell–cell junctions (Fig. S13). Moreover, this loss-and-regain of cell membrane probes can be repeated for at least 5 cycles without significant reduction in the efficiency (Fig. S13 and S14). As a result, the DNAMeter can now be used to study long-term cellular events.

Finally, we applied the EC22-DNAMeter to measure intercellular E-cadherin tensions during collective epithelial migrations. A slab of polydimethylsiloxane (PDMS) was pre-attached onto a substrate, and then a confluent MDCK cell monolayer was formed adjacent to the PDMS slab^36^. The interface between the monolayer and PDMS was defined as the initial edge. After the removal of the PDMS slab, the exposed free space triggers the migration of the cell sheet, emulating the wound healing process. After the initial force mapping with the EC22-DNAMeter, we replaced the HEPES-buffered saline with the complete cell growth medium. Following another 12 h of cell growth and migration, fresh EC22-DNAMeter was added to measure intercellular E-cadherin tensions (Fig. 5a and Fig. S15). Before removing the PDMS slab, intercellular forces mediated by E-cadherin were rarely observed within the cell sheet (Fig. 5b). After allowing the cell sheet to migrate for 12 h, interestingly, junctional pixels of high G/Y ratio were clearly observed in the regions ~15 cell lengths from the leading edge of migration (Fig. 5b). We have further quantified the correlation between the number of pixels experiencing >4.4 pN forces and their distance to the leading edge (Methods, Fig. S16 and Fig. 5c). As the distance increased, the percentage of E-cadherins undergoing intercellular tensions also linearly increased. In comparison, negligible forces were observed throughout the imaging zone before removing the PDMS (Fig. 5c). Overall, these observations are in good agreement with some previous studies on the global force distributions during this process^17,36,61^.

**Fig. 5.**
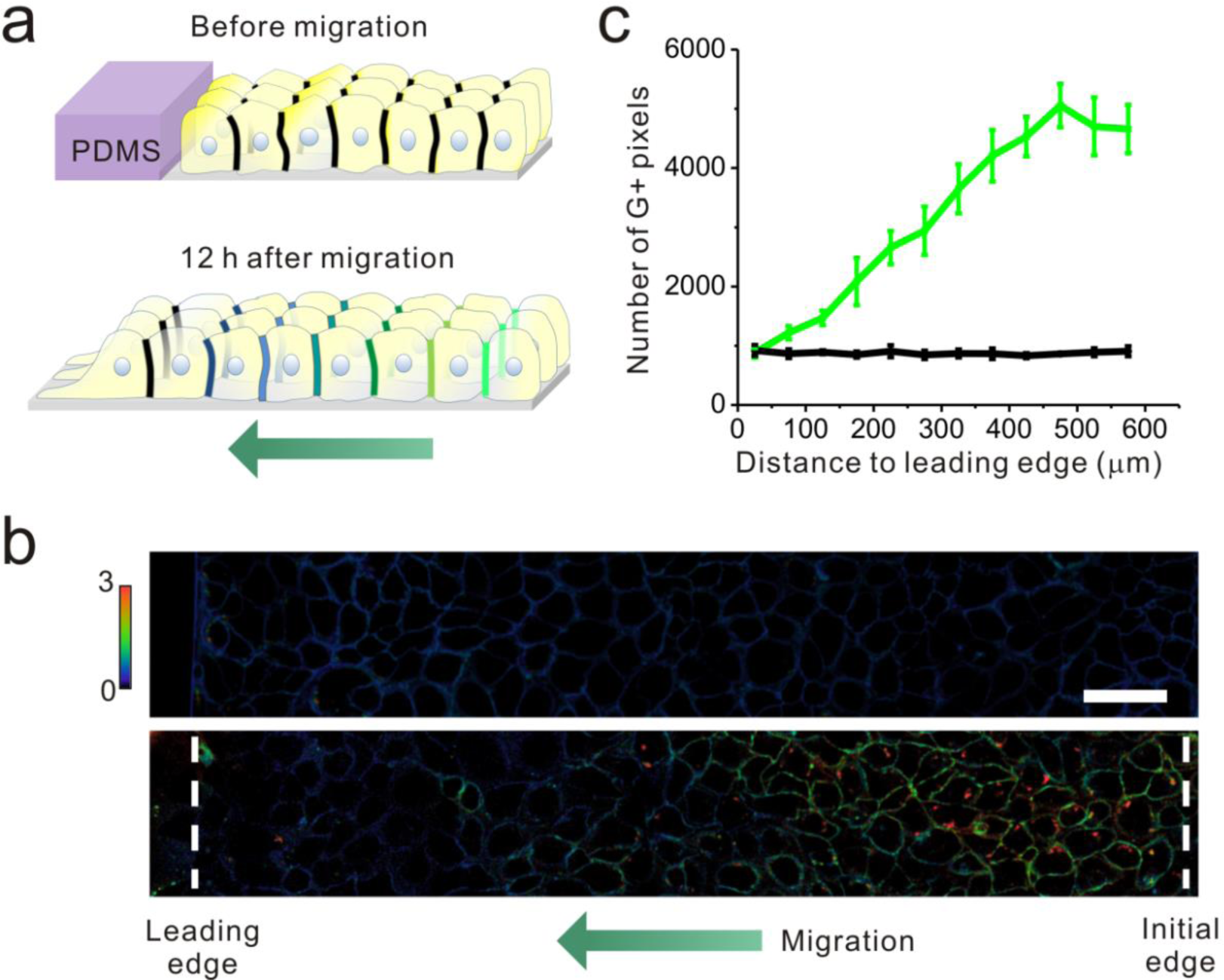
Mapping tensile force distributions during collective cell migration. (**a**) Schematic of collective MDCK cell migration. A PDMS slab was pre-attached on a glass bottom dish, and a confluent cell monolayer was formed next to it. Before removing the PDMS, the EC-DNAMeter was added to map the intercellular forces. The removal of the PDMS then triggered the collective migration. After 12 h of migration, fresh EC-DNAMeter was added again to map the forces. (**b**) Fluorescence imaging of MDCK monolayer cells before (top) and after 12 h of cell migration (bottom). G/Y ratio was shown and used for the quantification. Initial edge is the initial interface between the PDMS and monolayer cells before migration. Leading edge is the edge where “leader cells” located at the front edge of the advancing cell sheet. Scale bar, 50 µm. (**c**) Quantitative analysis of tension within these monolayer cells as a function of their distances to the leading edge before cell migration (black line) or after 12 h of cell migration (green line).

## CONCLUSIONS

In this study, we have developed a DNA-based probe to quantify, at the molecular level, E-cadherin-mediated tensile forces at cell–cell junctions. The so-called DNAMeter exhibits several unique features. First, the intrinsic modularity and precise self-assembly of the DNA scaffold allows the accurate positioning of specific reference fluorophores, reporter fluorophores and quenchers^32,48–50^. As a result, a facile ratiometric quantification of tensile forces can be achieved. Predictably, through the rational tuning of the sequence and length of the DNA hairpin, the threshold force can be tailored in a large range to study different types of intercellular forces^31,32^. By further conjugating two hairpins into one self-assembled “rod”-like DNA structure, a large range of tensile forces can be measured simultaneously. Compared to two separated membrane DNA hairpin probes, the conjugated DNAMeter allows the use of one reference fluorophore to characterize the membrane distributions of both hairpins. Supposedly, more hairpins can be incorporated into the DNAMeter to realize delicate quantification of an even larger range of forces.

Compared to the traction force microscopy^23,54^, the beauty of the DNAMeter is its capability to distinguish tension mediated by a particular protein from the total forces at cell–cell junctions. Compatible with readily accessible fluorescence microscopes, the DNAMeter is also easy to prepare and use. By simply incubating with the target cells, the DNAMeter can be spontaneously anchored onto cell membrane to report the tensions. Compared to fluorescent protein-based sensors and cantilever pillars^13,24^, there is no requirement for the cloning or microfabrication. In addition, the obtained fluorescence signals can be straightforwardly converted into mechanical forces without the need of complicated data processing or analysis. As a natural component in the cell plasma membrane, the cholesterol anchors can freely diffuse along the membrane^29^. In addition, the cholesterol-DNA conjugates can also be effectively removed if desired (Fig. S13 and S14).

We have demonstrated in detail two approaches to quantify the force distributions by either the number of bright pixels or the unfolded DNA probes. Both approaches can be facilely applied for mapping intercellular forces. In addition, the EC-DNAMeter has been used to quantify intercellular E-cadherin tension during the collective migration of cell sheet. In principle, the DNAMeter can also be used to quantify three-dimensional protein-specific intercellular forces in physiologically relevant multi-layer cell assemblies or tissues^51,52^. Our study demonstrated the ability of the DNAMeter to quantify and real-time monitor mechanical forces within a colony of cells. With a broad choice of fluorophores and quenchers, the DNAMeter can be further used to simultaneously measure intercellular forces among different receptor-ligand pairs, and to study the correlations between forces and the concentration gradients of morphogens or signaling molecules^53^. Further applications of the DNAMeter will allow the construction of more accurate mechanical models to study mechanotransduction during embryogenesis, morphogenesis, and various physiological and pathological processes.

## EXPERIMENTAL PROCEDURES

### Synthesis of Protein G-modified DNA strands

All the oligonucleotides were custom synthesized and purified by the W. M. Keck Oligonucleotide Synthesis Facility, unless otherwise noted. The sequences of these oligonucleotides were listed in Supplementary Table 1. For the synthesis of Protein G-modified DNA strands, 25 µL of thiol- and QSY^®^21-modified DNA ligand strand (200 µM) was first mixed with 10 µL of 100 mM TCEP in PBS buffer containing 50 mM EDTA at pH 7.2. After 1 h room temperature incubation to reduce the disulfide bonds, excess TCEP was removed using a Bio-Spin-6 column, followed by an immediate addition of 1.5 µL, 23 mM freshly prepared sulfo-SMCC. The mixture was then briefly incubated at room temperature for one min, and afterwards, 10 µL of 10 mg/mL Protein G was added. After an overnight incubation, the Protein G-DNA conjugates were purified with Dynabeads (Invitrogen) through a His-tag-specific purification. The final product was then buffer exchanged into DPBS, followed by concentrating and storage. The concentrations of Protein G-modified DNA strands were further quantified with a Nanodrop.

### Preparation of the EC-DNAMeter

To prepare the EC-DNAMeter, 1 µM of a cholesterol- and Cy5-modifed 22%GC hairpin, a FAM- and TAMRA-modified 66%GC hairpin, and a Dabcyl-labeled helper strand were first mixed at equal molar ratio in DPBS. After denaturing at 75°C for 5 min, the mixture was slowly annealed back to the room temperature at a rate of 1.3°C/min. These self-assembled DNAMeters were then incubated with an equal molar above-mentioned Protein G-modified DNA ligand strand for overnight reaction at 4°C. The as-prepared ProG-DNAMeter was then mixed with an equal molar IgG/Fc-fused Human E-cadherin (AcroBiosystems, catalog#: ECD-H5250) at room temperature for 15 min. The final EC-DNAMeter construct was thereby assembled and could be applied for the force measurement.

### F_1/2_ calculation for the DNA hairpin

F_1/2_ is defined as the force at which the DNA hairpin has 50% probability of being unfolded and can be calculated using the following equation^32^: F_1/2_= (ΔG_fold_+ΔG_stretch_) /Δx. Here, ΔG_fold_ is the free energy to unfold the DNA hairpin when no force is applied, which can be determined using nearest neighbor free energy parameters obtained from an IDT OligoAnalyzer software. ΔG_stretch_ is the free energy for stretching an unfolded single stranded DNA from no force up to F= F_1/2_. It can be calculated from a worm-like chain model^34^. Δx is the hairpin displacement length needed for unfolding and is estimated to be 0.44n+1.56 nanometer, where n is the length of the DNA hairpin, including both the stem and loop regions. The calculated F_1/2_ values for the 22%GC and 66%GC DNA hairpins were summarized in Table S2.

### In vitro fluorescence characterization of the probe

Fluorescence measurement was used to determine the sensitivity of the reporter fluorophore-quencher system and to validate the signal from the reference fluorophore. A dqEC-DNAMeter was prepared by adding to the EC-DNAMeter construct with excess amount of DNA strands that are complementary to the 22%GC and 66%GC hairpins. All the fluorescence measurements were performed with a PTI fluorimeter (Horiba, New Jersey, NJ). The excitation wavelength for the FAM, TAMRA and Cy5 was 488 nm, 557 nm and 640 nm, respectively, with a corresponding 510–560 nm, 560–610 nm and 650–700 nm emission spectra to be collected for each fluorophore.

### Preparation of the supported lipid monolayer

The supported lipid monolayers were prepared by adding a mixture of soybean polar extract and the DNAMeter onto Teflon AF-coated coverslips. A more detailed protocol was provided in our previous study^29^. Briefly, 1 µL of 1.2% Teflon AF solution, after diluting with Fluorinert FC-770, was added onto a clean coverslip and then spin coated at 2,000 rpm for 1 min. The coverslips were further dried at 180°C for 5 min to finish the coating. Afterwards, different concentrations of soybean polar extract/ DNAMeter mixture was added to form lipid monolayers.

### Calibration of the DNAMeter

To establish calibration curves to correlate the membrane fluorescence intensities with probe densities, different concentrations of DNAMeter were added onto the above-mentioned supported lipid monolayer. To prepare these DNAMeter-incorporated monolayers, soybean polar extract lipid solution was spiked with different concentrations of the DNA probe. After equilibrating at 4°C for overnight, 10 µL mixture was dried for 1 h under a reduced pressure to remove chloroform, and then rehydrated into 5 µL DPBS buffer. The obtained solution was then added on the above-prepared coverslips for the fluorescence imaging. The fluorescence intensity of the membrane DNAMeter was measured with a spinning disk confocal microscope, which data was further plotted as a function of the probe densities.

Here, the DNAMeter and dqDNAMeter was used as 0% unfolded and 100% unfolded probe, respectively. We mixed different amounts of the DNAMeter and dqDNAMeter in the supported lipid monolayer, and then imaged the corresponding membrane fluorescence of FAM (G, 485±10 nm excitation, 530±15 nm emission), TAMRA (Y, 540±20 nm excitation, 590±17 nm emission), and Cy5 (R, 624±20 nm excitation, 675±20 nm emission). The G/Y ratio of each individual pixel was used to calculate the percentage of unfolded 22%GC DNA hairpin, while the R/Y ratio was used for the 66%GC hairpin. A linear correlation was observed between the G/Y (or R/Y) ratio and the percentage of unfolded 22%GC (or 66%GC) DNA hairpin. This linear relationship was further used to convert the fluorescence signals on the cell membrane to the percentage of unfolded DNA probes.

### Cell culture and imaging

MDCK cells were cultured in DMEM medium supplemented with 10% FBS, 100 unit penicillin, and 0.1 mg/mL streptomycin. These cells were split at 80% confluency and plated at a density of 50% following standard cell culture procedures. All images were collected with an NIS-Elements AR software using a Yokogawa spinning disk confocal on a Nikon Eclipse-TI inverted microscope. FAM was excited with a 488 nm laser line. TAMRA and Cy5 were exited with 561 and 640 nm laser line, respectively. Data analysis was performed with an NIS-Elements AR Analysis software.

### Imaging of E-cadherin-mediated tensile forces

To directly image E-cadherin-mediated tension, MDCK cells were first seeded on a glass bottom dish and grown overnight. After washing twice with HEPES-buffered saline (Live Cell Imaging Solution, Invitrogen), 0.2 µM pre-assembled EC-DNAMeter was added. After room temperature incubation in the HEPES-buffered saline solution for 1 h, unbound EC-DNAMeter was washed away with HEPES-buffered saline for three times. Cell imaging was then followed immediately with a 100× oil immersion objective. To track the dynamics of EGTA-treated E-cadherin tension, after adding 0.2 µM EC-DNAMeter for 1 h and washing away unbound probes, 10 mM EGTA was added followed by immediate force imaging every 5 min for a total of 30 min.

### Determining the percentage of force-experiencing pixels at cell–cell junctions

For this purpose, cellular background fluorescence was first subtracted in each image. Ratiometric images were then generated by dividing the reporter fluorescence (G or R) with the reference fluorescence (Y) using an NIS-Elements AR Analysis software. We then generated a pixel distribution plot by pinning the number of pixels exhibiting similar range of G/Y or R/Y ratios versus the corresponding ratio. Based on a control ProG-DNAMeter, we could estimate a threshold value of G/Y (1.0) and R/Y (0.24) to distinguish force-experiencing pixels from the background. We could then count the number of pixels showing positive signals, i.e., G/Y in the range of 1.0–3.0 or R/Y in the range of 0.24–1.0, denoted as N_G+_ and N_G+R+_, respectively. The percentage of pixels involved in >8.1 pN force events was then calculated by dividing the number of pixels exhibiting both positive G/Y and positive R/Y, denoted as N_G+R+_, by the total pixel number N_0_. In other words, (N_G+R+_/N_0_) × 100%. Similarly, the percentage of pixels experiencing >4.4 pN forces was calculated by dividing G/Y-positive pixel number, N_G+_, by the total pixel number N_0_. In other words, (N_G+_/N_0_) × 100%. In this way, the percentage of pixels involved in 4.4–8.1 pN force events could also be calculated by subtracting (N_G+R+_/N_0_) × 100% from (N_G+_/N_0_) × 100%.

### Determining the percentage of unfolded probes at cell–cell junctions

To determine the percentage of unfolded probes at a cell–cell junction, pixels with distinguishable TAMRA fluorescence were first grouped into different subranges. The same subrange of pixels exhibiting similar level of G/Y or R/Y ratios. The interval between each subrange was determined based on the standard deviation of the fluorescence ratios, i.e., G/Y 0.25 and R/Y 0.12. Afterwards, the percentage of unfolded probes at each cell–cell junction was calculated by 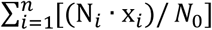. Here, N_*i*_ is the number of pixels in each subrange, and N_0_ is the total number of pixels at the junction. In this equation, x_*i*_ is the corresponding percentage of unfolded hairpin in each pixel subrange, which can be determined from the above-mentioned linear standard curve from the supported lipid monolayer measurement. The fraction of unfolded probes by >4.4 pN and >8.1 pN tension was thereby calculated based on the G/Y and R/Y ratio, respectively. The fraction of unfolded probe experiencing 4.4–8.1 pN forces was then calculated by subtracting the fraction of unfolded probes by >8.1 pN from that of probes by >4.4 pN tension.

### Collective cell migration

A slab of PDMS was gently pressed against the bottom of a glass bottom dish, followed by seeding the MDCK cells. After 48 h growth, the confluent cell monolayer was washed with HEPES-buffered saline for three times. After adding 0.2 µM EC22-DNAMeter at room temperature for 1 h, the probe-anchored MDCK cell monolayer was washed twice with HEPES-buffered saline, and a large area scan was immediately conducted with a 40× oil immersion objective. After imaging, probe-containing solution was discarded and replaced with the complete cell growth medium, followed by the removal of the PDMS slab. After another 12 h of cell growth, the medium was removed and the cell monolayer was washed with HEPES-buffered saline for three times, followed by the addition of 0.2 µM EC22-DNAMeter. After 1 h incubation at room temperature, the cell monolayer was washed twice with HEPES-buffered saline and imaged again in the large area mode with the confocal microscope.

We further quantitatively correlated the number of pixels involved in >4.4 pN tension with their distances to the leading edge. Here, we counted the number of pixels per unit area exhibiting high G/Y ratio. In total, 2,500 unit areas were selected continuously across the whole imaging area from the leading edge (Fig. S10). The number of pixels with positive ratios in the range of 1.0–3.0 was then counted and further plotted with their distances to the leading edge.

## ACKNOWLEDGEMENTS

The authors gratefully acknowledge NIH R35GM133507, a start-up grant from UMass Amherst and IALS M2M seed grant to M. You and NSF CMMI 1662835 to Y. Sun. We are grateful to Dr. James Chambers for the assistance in fluorescence imaging, and Dr. Tianxi Yang for manuscript preparation. We also thank every other member of the You Lab and Dr. Craig Martin for useful discussion.

## AUTHOR CONTRIBUTIONS

M.Y. provided the guidance for the whole project. B.Z. and M.Y. developed the experimental plan. B.Z. designed/synthesized DNA probes and performed all the cellular imaging and data analysis. N.L., B.Z., Y.S. and M.Y. designed the collective cell migration experiment and analyzed the data. T.X. and B.Z. performed western blot and immunofluorescence staining experiments. C.L. performed computational simulation and data analysis. Y.B. conducted cell culture and analyzed the data. B.Z. and M.Y. wrote the manuscript, and all other authors have reviewed and edited the manuscript.

## REFERENCES AND NOTES

1 Charras, G. & Yap, A. S. Tensile forces and mechanotransduction at cell–cell Junctions. Curr. Biol. 28, R445–R457 (2018).

2 Wozniak, M. A. & Chen, C. S. Mechanotransduction in development: a growing role for contractility. Nat. Rev. Mol. Cell Biol. 10, 34–43 (2009).

3 Chen, C. S., Tan, J. & Tien, J. Mechanotransduction at cell-matrix and cell–cell contacts. Annu. Rev. Biomed. Eng. 6, 275–302 (2004).

4 Levine, E., Lee, C. H., Kintner, C. & Gumbiner, B. M. Selective disruption of E-cadherin function in early Xenopus embryos by a dominant negative mutant. Development 120, 901–909 (1994).

5 Martin, A. C., Kaschube, M. & Wieschaus, E. F. Pulsed contractions of an actin–myosin network drive apical constriction. Nature 457, 495 (2008).

6 Bazellières, E. et al. Control of cell–cell forces and collective cell dynamics by the intercellular adhesome. Nat. Cell Biol. 17, 409–420 (2015).

7 Acharya, B. R. et al. Mammalian Diaphanous 1 mediates a pathway for E-cadherin to stabilize epithelial barriers through junctional contractility. Cell Rep. 18, 2854–2867 (2017).

8 Conway, Daniel E. et al. Fluid shear stress on endothelial cells modulates mechanical tension across VE-Cadherin and PECAM-1. Curr. Biol. 23, 1024–1030 (2013).

9 Takeichi, M. The cadherins: cell–cell adhesion molecules controlling animal morphogenesis. Development 102, 639–655 (1988).

10 van Roy, F. & Berx, G. The cell–cell adhesion molecule E-cadherin. Cell. Mol. Life Sci. 65, 3756–3788 (2008).

11 Leckband, D. E., le Duc, Q., Wang, N. & de Rooij, J. Mechanotransduction at cadherin-mediated adhesions. Curr. Opin. Cell Biol. 23, 523–530 (2011).

12 Niessen, C. M., Leckband, D. & Yap, A. S. Tissue organization by cadherin adhesion molecules: dynamic molecular and cellular mechanisms of morphogenetic regulation. Physiol. Rev. 91, 691–731 (2011).

13 Liu, Z. et al. Mechanical tugging force regulates the size of cell–cell junctions. Proc. Natl. Acad. Sci. U. S. A. 107, 9944–9949 (2010).

14 Halbleib, J. M. & Nelson, W. J. Cadherins in development: cell adhesion, sorting, and tissue morphogenesis. Genes Dev. 20, 3199–3214 (2006).

15 Stemmler, M. P. Cadherins in development and cancer. Mol. BioSyst. 4, 835–850 (2008).

16 Brugués, A. et al. Forces driving epithelial wound healing. Nat. Phys. 10, 683–690 (2014).

17 Trepat, X. et al. Physical forces during collective cell migration. Nat. Phys. 5, 426–430 (2009).

18 van Roy, F. Beyond E-cadherin: roles of other cadherin superfamily members in cancer. Nat. Rev. Cancer 14, 121–134 (2014).

19 Leckband, D. E. & Rooij, J. d. Cadherin adhesion and mechanotransduction. Annu. Rev. Cell Dev. Biol. 30, 291–315 (2014).

20 Polacheck, W. J. & Chen, C. S. Measuring cell-generated forces: a guide to the available tools. Nat. Methods 13, 415–423 (2016).

21 Roca-Cusachs, P., Conte, V. & Trepat, X. Quantifying forces in cell biology. Nat. Cell Biol. 19, 742–751 (2017).

22 Eisenstein, M. Mechanobiology: a measure of molecular muscle. Nature 544, 255–257 (2017).

23 Tambe, D. T. et al. Collective cell guidance by cooperative intercellular forces. Nat. Mater. 10, 469–475 (2011).

24 Borghi, N. et al. E-cadherin is under constitutive actomyosin-generated tension that is increased at cell–cell contacts upon externally applied stretch. Proc. Natl. Acad. Sci. U. S. A. 109, 12568–12573 (2012).

25 Hoffman, B. D. & Yap, A. S. Towards a dynamic understanding of cadherin-based mechanobiology. Trends Cell Biol. 25, 803–814 (2015).

26 Wang, P. et al. Visualizing spatiotemporal dynamics of intercellular mechanotransmission upon wounding. ACS Photonics 5, 3565–3574 (2018).

27 Jurchenko, C. & Salaita, K. S. Lighting up the force: investigating mechanisms of mechanotransduction using fluorescent tension probes. Mol. Cell. Biol. 35, 2570–2582 (2015).

28 Zhao, B. et al. Visualizing intercellular tensile forces by DNA-based membrane molecular probes. J. Am. Chem. Soc. 139, 18182–18185 (2017).

29 You, M. X. et al. DNA probes for monitoring dynamic and transient molecular encounters on live cell membranes. Nat. Nanotechnol. 12, 453–459 (2017).

30 Bunge, A. et al. Lipid membranes carrying lipophilic cholesterol-based oligonucleotides—characterization and application on layer-by-layer coated particles. J. Phys. Chem. B 113, 16425–16434 (2009).

31 Wang, X. & Ha, T. Defining single molecular forces required to activate integrin and notch signaling. Science 340, 991–994 (2013).

32 Zhang, Y., Ge, C., Zhu, C. & Salaita, K. DNA-based digital tension probes reveal integrin forces during early cell adhesion. Nat. Commun. 5, 5167 (2014).

33 Blakely, B. L. et al. A DNA-based molecular probe for optically reporting cellular traction forces. Nat. Methods 11, 1229–1232 (2014).

34 Woodside, M. T. et al. Nanomechanical measurements of the sequence-dependent folding landscapes of single nucleic acid hairpins. Proc. Natl. Acad. Sci. U. S. A. 103, 6190–6195 (2006).

35 Kronenberg, N. M. et al. Long-term imaging of cellular forces with high precision by elastic resonator interference stress microscopy. Nat. Cell Biol. 19, 864–872 (2017).

36 Vedula, S. R. K. et al. Emerging modes of collective cell migration induced by geometrical constraints. Proc. Natl. Acad. Sci. U. S. A. 109, 12974–12979 (2012).

37 Li, L., He, Y., Zhao, M. & Jiang, J. Collective cell migration: implications for wound healing and cancer invasion. Burns & Trauma 1, 21–26 (2013).

38 Marko, J. F. & Siggia, E. D. Stretching DNA. Macromolecules 28, 8759–8770 (1995).

39 Smith, S. B., Cui, Y. & Bustamante, C. Overstretching B-DNA: the elastic response of individual double-stranded and single-stranded DNA molecules. Science 271, 795–799 (1996).

40 Wang, X. et al. Constructing modular and universal single molecule tension sensor using protein G to study mechano-sensitive receptors. Sci. Rep. 6, 21584 (2016).

41 Liu, H. et al. DNA-based micelles: synthesis, micellar properties and size-dependent cell permeability. Chem. Eur. J. 16, 3791–3797 (2010).

42 Stetter, F. W. S., Cwiklik, L., Jungwirth, P. & Hugel, T. Single lipid extraction: the anchoring strength of cholesterol in liquid-ordered and liquid-disordered phases. Biophys. J. 107, 1167–1175 (2014).

43 Hannestad, J. K. et al. Kinetics of diffusion-mediated DNA hybridization in lipid monolayer films determined by single-molecule fluorescence spectroscopy. ACS Nano 7, 308–315 (2013).

44 Muhamed, I. et al. E-cadherin-mediated force transduction signals regulate global cell mechanics. J. Cell Sci. 129, 1843–1854 (2016).

45 Pokutta, S., Herrenknecht, K., Kemler, R. & Engel, J. Conformational changes of the recombinant extracellular domain of E-cadherin upon calcium binding. Eur. J. Biochem. 223, 1019–1026 (1994).

46 Rothen-Rutishauser, B., Riesen, F. K., Braun, A., Günthert, M. & Wunderli-Allenspach, H. Dynamics of tight and adherens junctions under EGTA treatment. J. Membr. Biol. 188, 151–162 (2002).

47 Pandya, P., Orgaz, J. L. & Sanz-Moreno, V. Actomyosin contractility and collective migration: may the force be with you. Curr. Opin. Cell Biol. 48, 87–96 (2017).

48 Surana, S., Bhat, J. M., Koushika, S. P. & Krishnan, Y. An autonomous DNA nanomachine maps spatiotemporal pH changes in a multicellular living organism. Nat. Commun. 2, 340 (2011).

49 Chakraborty, K., Veetil, A. T., Jaffrey, S. R. & Krishnan, Y. Nucleic acid–based nanodevices in biological imaging. Annu. Rev. Biochem. 85, 349–373 (2016).

50 Ranallo, S., Porchetta, A. & Ricci, F. DNA-based scaffolds for sensing applications. Anal. Chem. 91, 44–59 (2019).

51 Legant, W. R. et al. Microfabricated tissue gauges to measure and manipulate forces from 3D microtissues. Proc. Natl. Acad. Sci. U. S. A. 106, 10097–10102 (2009).

52 Nelson, C. M. From static to animated: measuring mechanical forces in tissues. J. Cell Biol. 216, 29–30 (2017).

53 Tabata, T. & Takei, Y. Morphogens, their identification and regulation. Development 131, 703–712 (2004).

54 Maruthamuthu, V., Sabass, B., Schwarz, U. S. & Gardel, M. L. Cell-ECM traction force modulates endogenous tension at cell-cell contacts. Proc. Natl. Acad. Sci. U. S. A. 108, 4708–4713 (2011).

55 Bagheri, Y., Chedid, S., Shafiei, F., Zhao, B. & You, M. A quantitative assessment of the dynamic modification of lipid-DNA probes on live cell membranes. Chem. Sci. 10, 11030–11040 (2019).

56 Twiss, F. et al. Vinculin-dependent cadherin mechanosensing regulates efficient epithelial barrier formation. Biol. Open 1, 1128–1140 (2012).

57 Seddiki, R. et al. Force-dependent binding of vinculin to α-catenin regulates cell-cell contact stability and collective cell behavior. Mol. Biol. Cell 29, 380–388 (2018).

58 Tian, X. et al. E-cadherin/β-catenin complex and the epithelial barrier. J. Biomed. Biotechnol. 567305 (2011).

59 Hartsock, A. & Nelson, W. J. Adherens and tight junctions: structure, function and connections to the actin cytoskeleton. Biochim. Biophys. Acta. 1778, 660–669 (2008).

60 Shewan, A. M., Maddugoda, M., Kraemer, A., Stehbens, S. J., Verma, S., Kovacs, E. M. & Yap, A. S. Myosin 2 is a key Rho kinase target necessary for the local concentration of E-cadherin at cell-cell contacts. Mol. Biol. Cell 16, 4531–4542 (2005).

61 Gayrard, C., Bernaudin, C., Dejardin, T., Seiler, C. & Borghi, N. Src- and confinement-dependent FAK activation causes E-cadherin relexation and β-catenin activity. J. Cell Biol. 217, 1063–1077.

