## Supplemental Information for "Quantifying Tensile Forces at Cell–Cell Junctions with a DNA-based Fluorescent Probe"

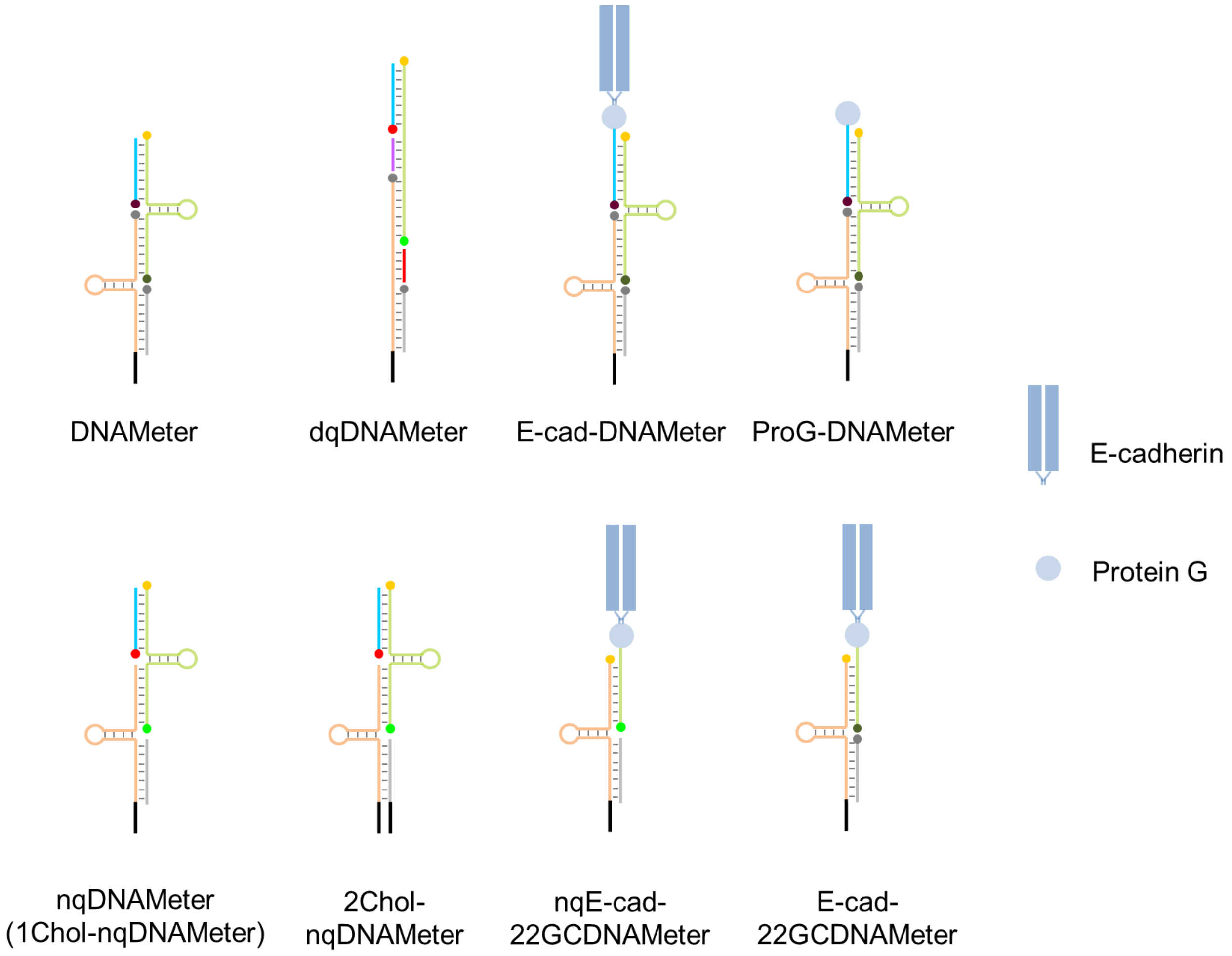

**Supplementary Scheme 1.** Schematic of different versions of DNAMeters used in this project.

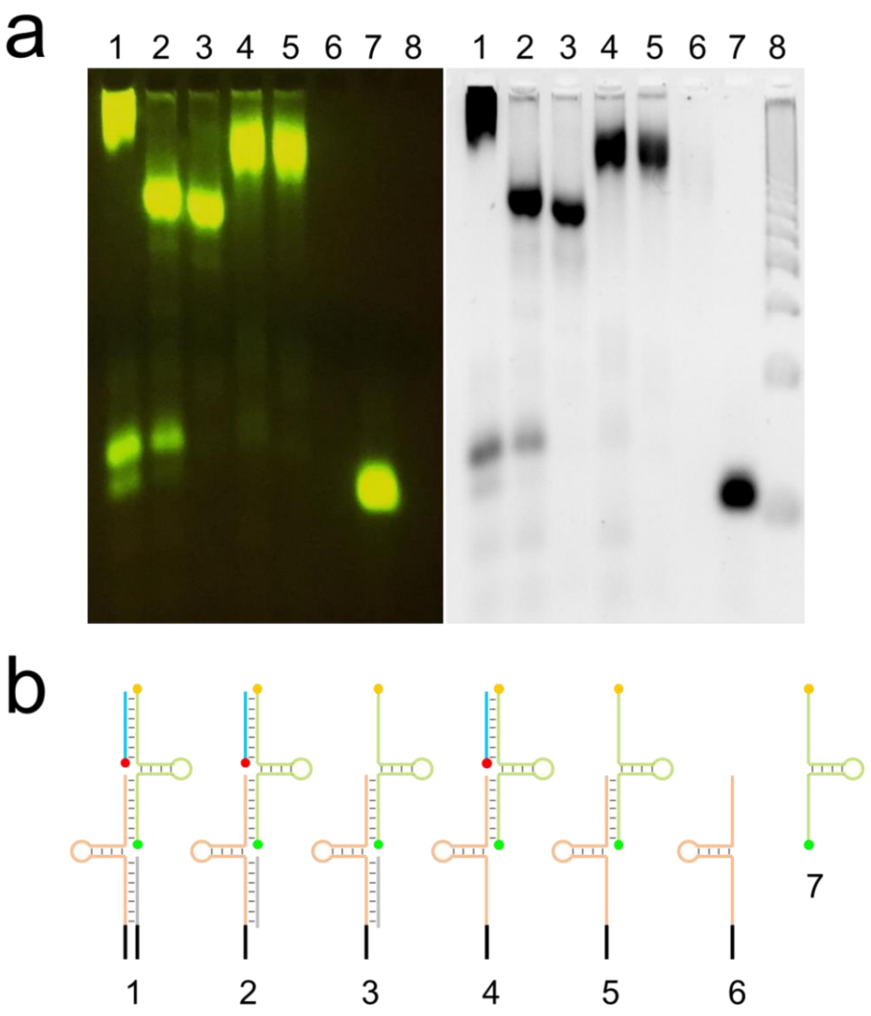

**Supplementary Figure 1. Gel mobility shift assay for the characterization of the nqDNAMeters.**

(**a**) 4% agarose gel electrophoresis before (left) and after (right) SYBR safe staining. The structure and DNA strand composition of each lane is shown in the panel (b). Lane 8 is a 50 bp DNA ladder (NEB). With SYBR safe staining, intense bands were observed because SYBR safe has similar excitation window as that of FAM.

(**b**) Schematics of the DNA structures in Lane 1–7. The green strand represents a TAMRA- and FAM-labeled 66%GC hairpin. The orange strand represents a 22%GC hairpin. The grey strand is a helper strand. The blue one is a Cy5-labeled ligand strand. Each black line represents a cholesterol anchor.

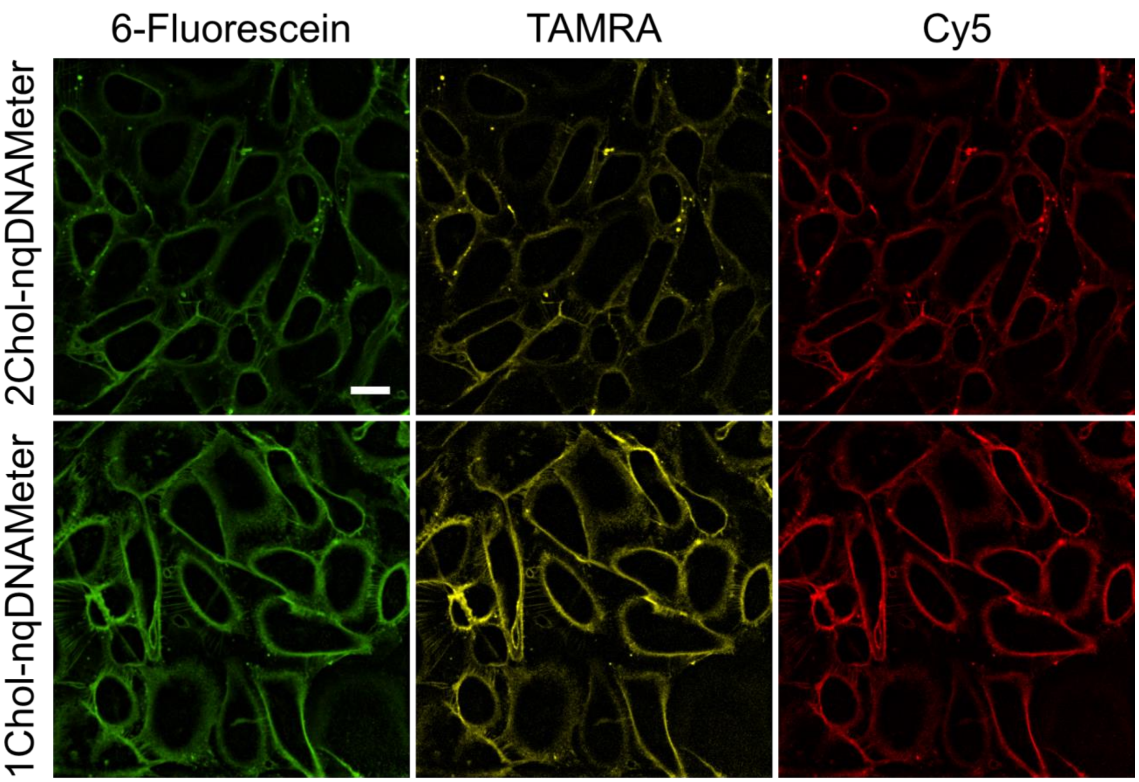

**Supplementary Figure 2. The insertion efficiency of one or two cholesterol-modified DNAMeter on live MDCK cell membrane.**

In this experiment, 1 µM of 1Chol-nqDNAMeter or 2Chol-nqDNAMeter was incubated with MDCK cells at room temperature for 30 min. A representative cell region was shown for each fluorescence channel by imaging with a spinning disk confocal microscope with a 40× oil immersion objective. Scale bar, 20 µm. Our results showed that 1Chol-nqDNAMeter exhibited brighter membrane fluorescence signals on MDCK cells.

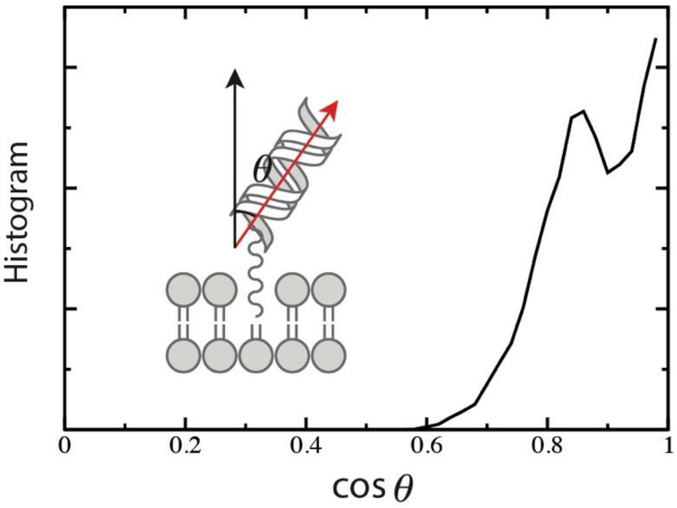

**Supplementary Figure 3. The distribution of the DNAMeter tilting angle with respect to the membrane surface normal.**

**Supplementary Note 1. Molecular dynamics simulation procedure**

Classical molecular dynamics simulations using atomistic models were performed using a GROMACS 2018 package [1]. A CHARMM36 [2] force field was chosen for modeling DNA and lipid molecules with a TIP3P water model [3]. First, the lipid bilayer of mixed DOPC and DOPG was constructed using a Charmm-GUI web server [4] containing 50 DOPC and 50 DOPG. The membrane normal was aligned with the Z direction. The initial DNA configuration was generated using a DNA builder web server [5] to adopt the form of B-DNA. Then, the DNA probe (5'-Cholesterol-C3-T21-3' / 5'-A21-3') was placed in the center of the simulation box with the size of $5.85 \times 5.85 \times 19\ nm^3$. The Z position of the cholesterol was aligned with that of the hydrophobic lipid tail in the upper membrane leaflet. The system was then solvated by ~17,000 water molecules with 140 mM NaCl to reproduce the experimental conditions. After steepest descent energy minimization, an equilibration simulation was run at a constant temperature (310 K) and pressure (1 atm), both coupled with the Berendsen method [6] for 100 ns. Then 100 ns NVT simulation was performed using a velocity rescaling method [7] with an external heat bath at 300 K (coupling time 1 ps). After that, the system was equilibrated. During the production runs, the LINCS algorithm [8] was used to constrain bond lengths and angles of the protein, allowing an integration time step of 2 fs. Long-range electrostatic interactions beyond a cutoff of 1.2 nm were calculated by the Particle-Mesh-Ewald (PME) method [9] with a grid spacing of 0.12 nm. Short-range repulsive and attractive dispersion interactions were described with the Lennard-Jones potentials, using 1.2 nm for the cutoff length. The temperature of each replica was then controlled using a velocity-rescaling method [7] with an external heat bath at target temperature with the coupling time of 1 ps. The volume of the simulation boxes was kept constant. The total sampling time of the production run was 1 μs.

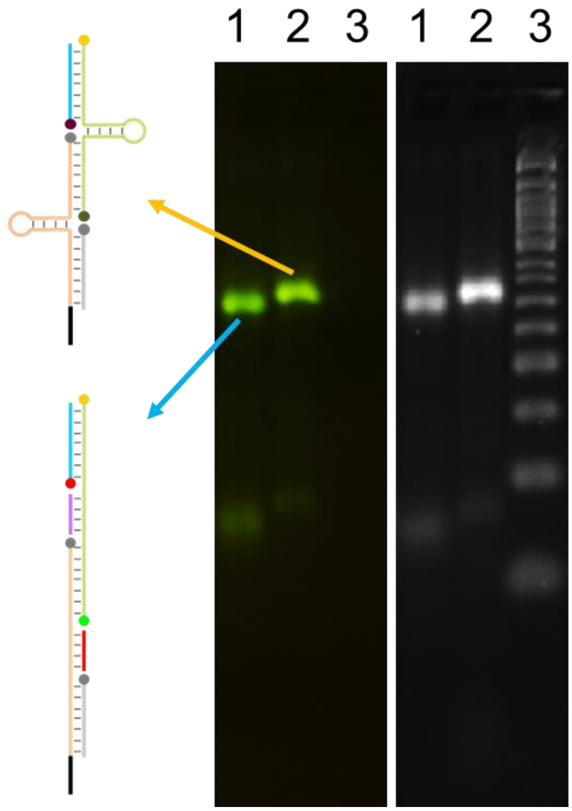

**Supplementary Figure 4. Gel mobility shift assay for the DNAMeter and dqDNAMeter before (left) and after (right) SYBR safe staining.**

Schematics of the corresponding DNA structures have been shown. The green strand represents a TAMRA- and FAM-labeled 66%GC hairpin. The orange strand represents a 22%GC hairpin. The grey strand is a Dabcyl-labeled helper strand. The blue one is a Cy5-labeled ligand strand. The black line represents a cholesterol anchor. The red and purple strands are the DNA strands that are complementary to the 22%GC and 66%GC hairpin, respectively. Lane 3 is a 50 bp DNA ladder (NEB).

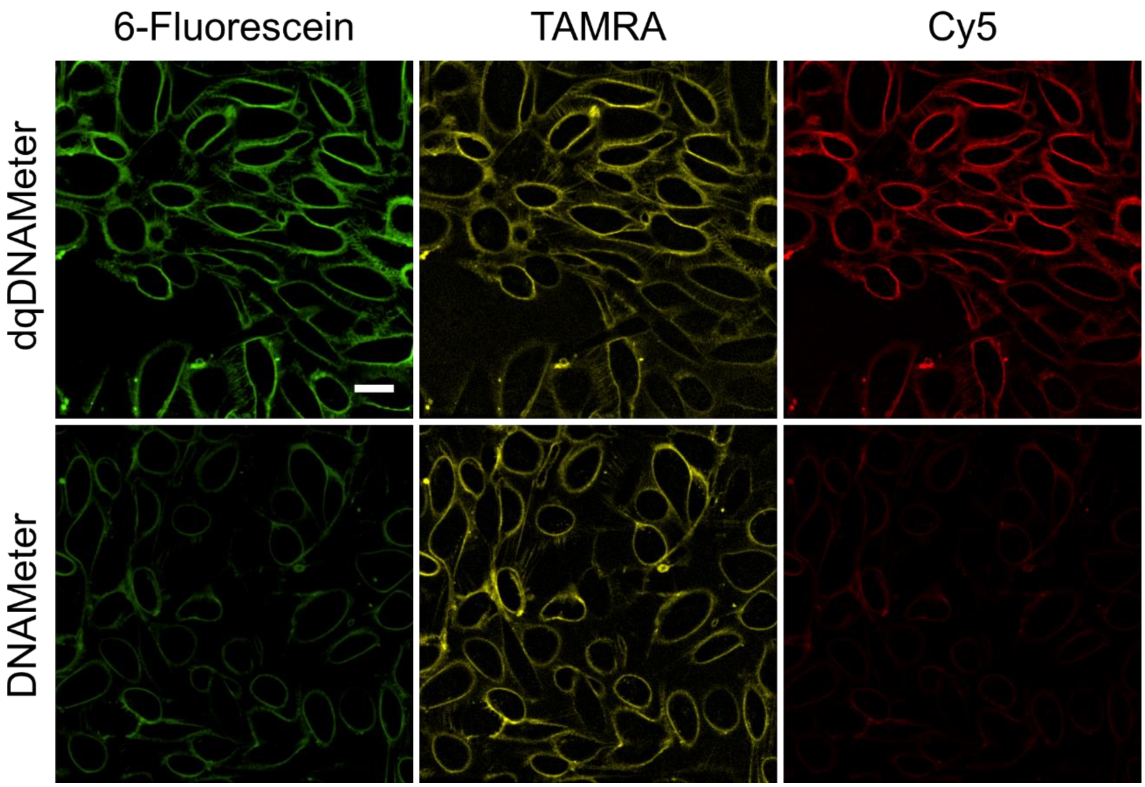

**Supplementary Figure 5. Membrane anchoring and performance of the DNAMeter and dqDNAMeter on live MDCK cells.**

In this experiment, 1 µM of DNAMeter or dqDNAMeter was incubated with MDCK cells at room temperature for 30 min. A representative cell region was shown for each fluorescence channel using a spinning disk confocal microscope with a 40× oil immersion objective. Scale bar, 20 µm.

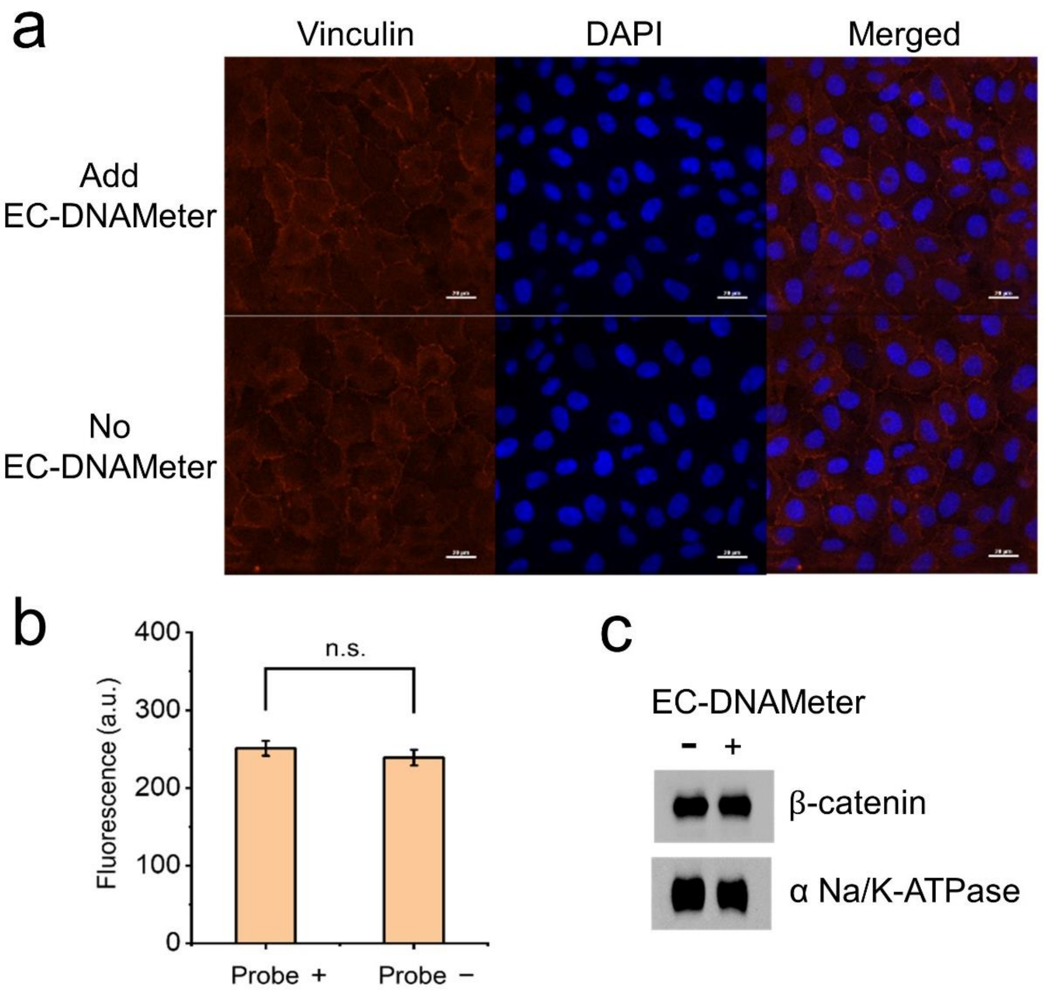

**Supplementary Figure 6. Studying the effect of membrane anchoring of the EC-DNAMeter on the adhesion and mechanical functions of MDCK cells.**

(**a**) Immunofluorescence staining of vinculin in MDCK cells with or without adding EC-DNAMeter. DAPI was used to stain the nucleus of the MDCK cells. Scale bar, 20 µm.

(**b**) Quantitative comparison of vinculin fluorescence intensities at different MDCK cell–cell junctions (N= 50), in the presence (Probe +) or absence (Probe -) of EC-DNAMeter. No significant difference in the fluorescence intensities were observed ($p > 0.05$).

(**c**) Western blot analysis of β-catenin isolated from plasma membrane of MDCK cells in the presence or absence of EC-DNAMeter. Here, α-Na/K ATPase was used as the loading control for normalizing the isolated membrane proteins.

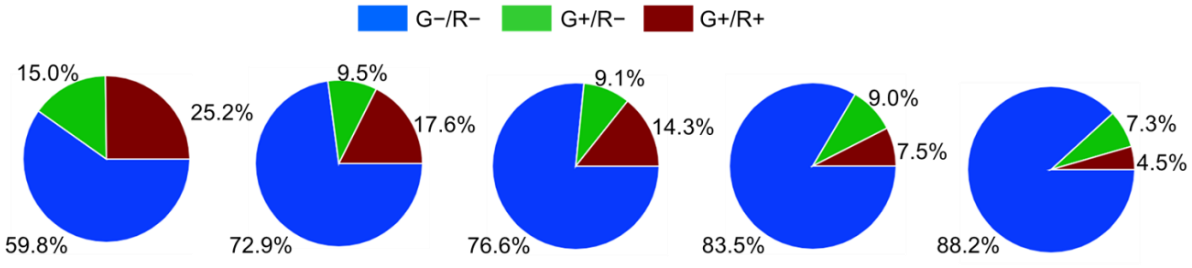

**Supplementary Figure 7. EGTA-induced dynamic disruption of intercellular forces.**

Fluorescence imaging of a representative cell–cell junction upon the EGTA treatment have been shown in Figure 4a. Here we illustrated the quantitative analysis of the percentage of pixels experiencing tensile forces at this cell–cell junction. The blue, green, and red region indicted the distribution of tensile forces in the range of below 4.4 pN, 4.4–8.1 pN, and above 8.1 pN, respectively.

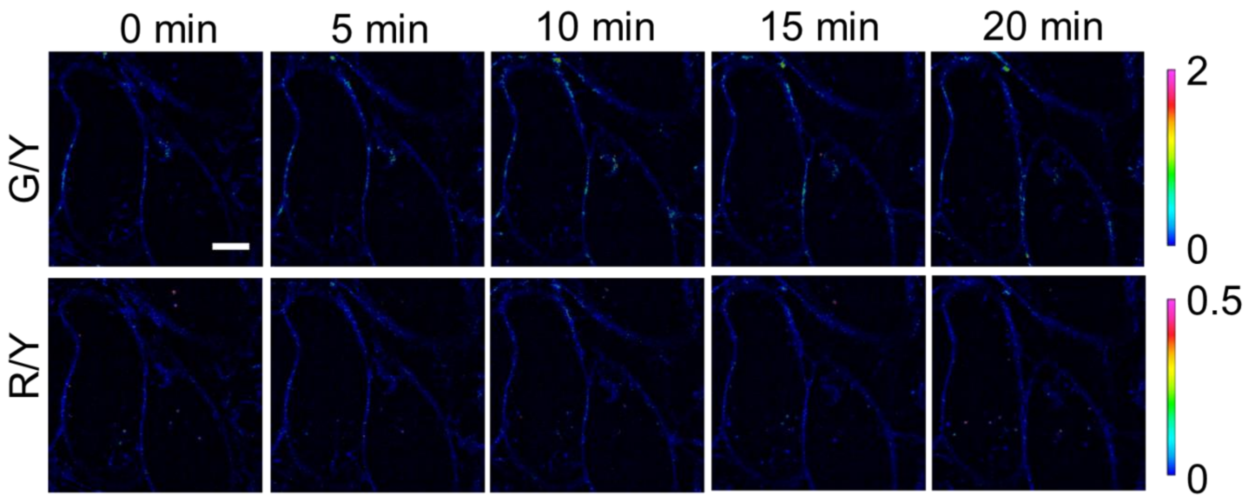

**Supplementary Figure 8. Ratiometric imaging of ProG-DNAMeter-modified MDCK cells after the addition of EGTA.**

At 0 min, 10 mM EGTA was added. G/Y stands for the ratio of green fluorescence (FAM) to yellow fluorescence (TAMRA). R/Y stands for red (Cy5)-to-yellow (TAMRA) fluorescence ratio. Scale bar, 5 µm.

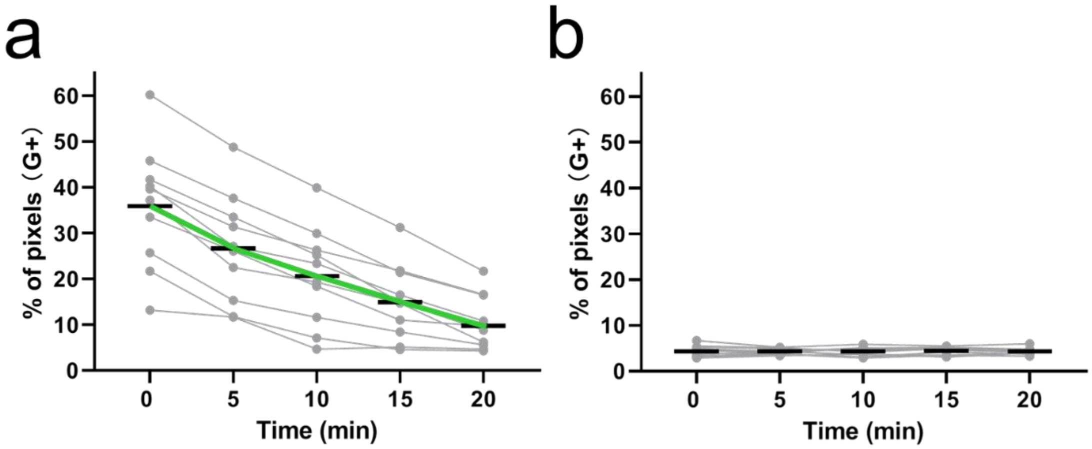

**Supplementary Figure 9. Statistical analysis of the dynamic changes in the percentage of pixels experiencing tension upon adding EGTA.**

Correlated with the data shown in Figure 4c and 4d, we illustrated here the percentage of pixels experiencing >4.4 pN tension (G+) at MDCK cell–cell junctions upon adding 10 mM EGTA (N= 10) at 0 min using the (**a**) EC-DNAMeter or (**b**) ProG-DNAMeter.

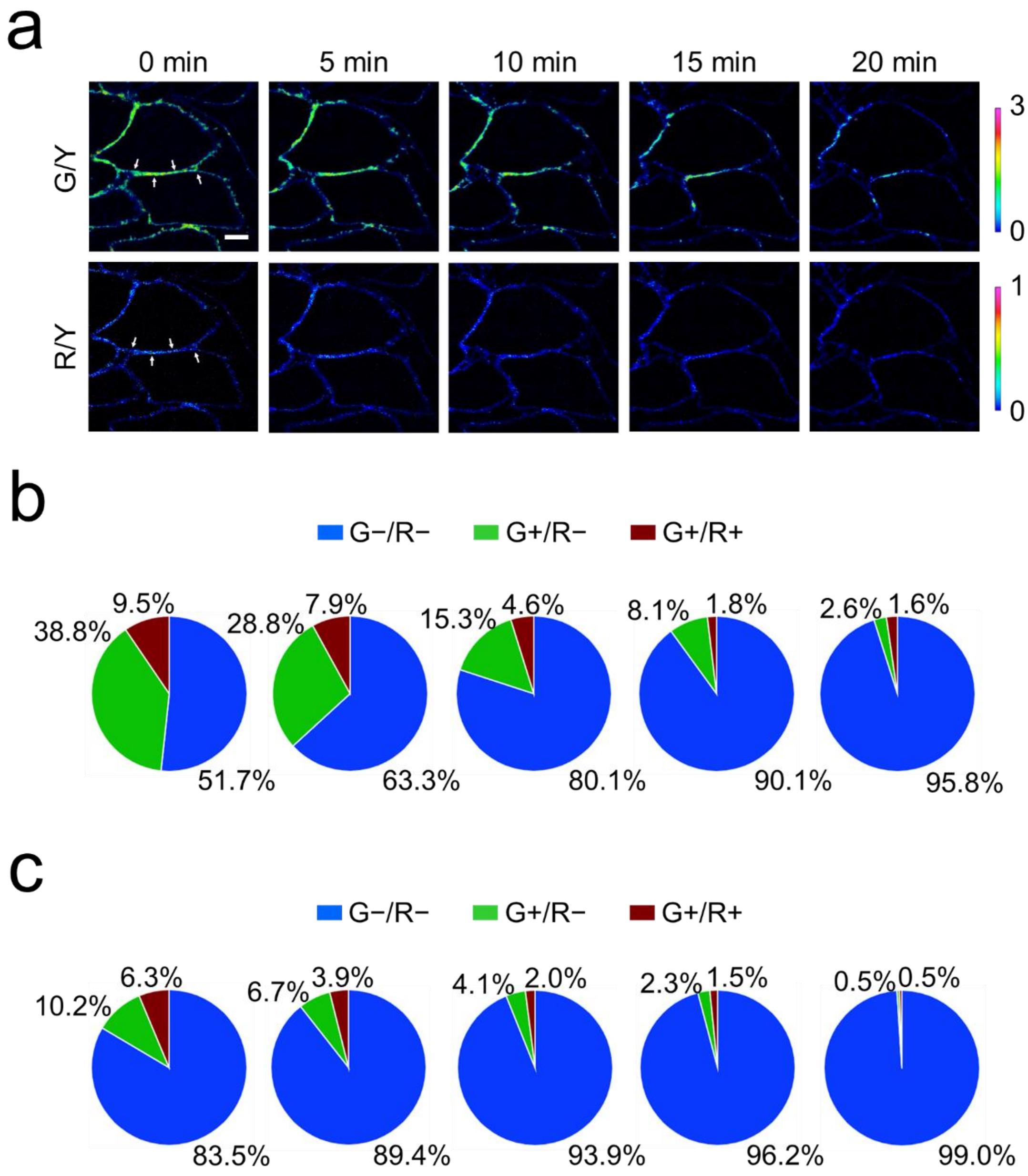

**Supplementary Figure 10. Monitoring ML-7-induced changes in E-cadherin tensions.**

(**a**) Representative ratiometric images of EC-DNAMeter-modified MDCK cells after adding 100 µM ML-7 at 0 min. The cell–cell junction denoted by white arrows was used for the quantitative analysis in the panel (b) and (c). Scale bar, 5 µm.

(**b**) The quantitative analysis of tension revealed by the percentage of pixels experiencing forces at different time after adding ML-7, as quantified with the EC-DNAMeter. Each pie chart corresponds to the image above it in the panel (a). The blue, green, and red region indicted the distribution of tensile forces in the range of <4.4 pN, 4.4–8.1 pN, and >8.1 pN, respectively.

(**c**) The quantitative analysis of tension revealed by the percentage of unfolded EC-DNAMeter probes at different time after adding ML-7. Each pie chart corresponds to the image above it in the panel (a). The blue, green, and red region indicted the distribution of tensile forces in the range of <4.4 pN, 4.4–8.1 pN, and >8.1 pN, respectively.

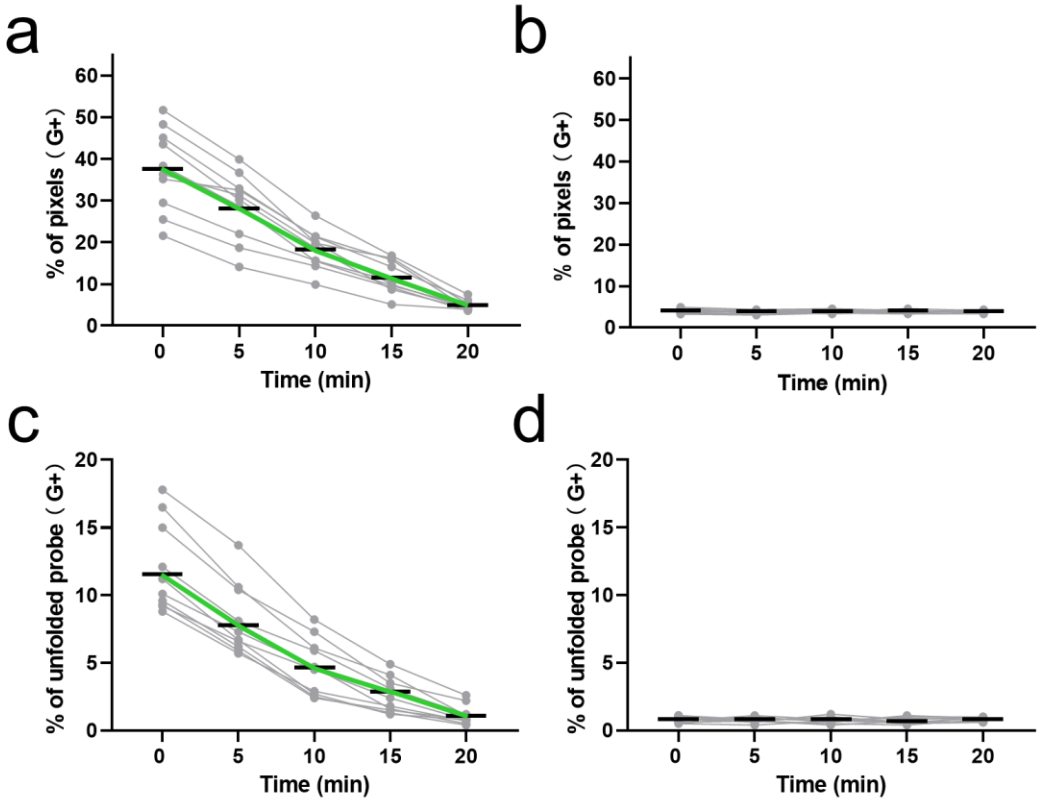

**Supplementary Figure 11. Statistical analysis of the dynamics of ML-7-induced changes in E-cadherin tensions in MDCK cells.**

(**a, b**) Statistical analysis of the dynamic changes in the percentage of pixels experiencing tension (>4.4 pN) with the (a) EC-DNAMeter or (b) ProG-DNAMeter, at different time after adding 100 µM ML-7. N= 10.

(**c, d**) Statistical analysis of the dynamic changes in the percentage of unfolded probes experiencing tension (>4.4 pN) with the (c) EC-DNAMeter or (d) ProG-DNAMeter, at different time after adding 100 µM ML-7. N= 10.

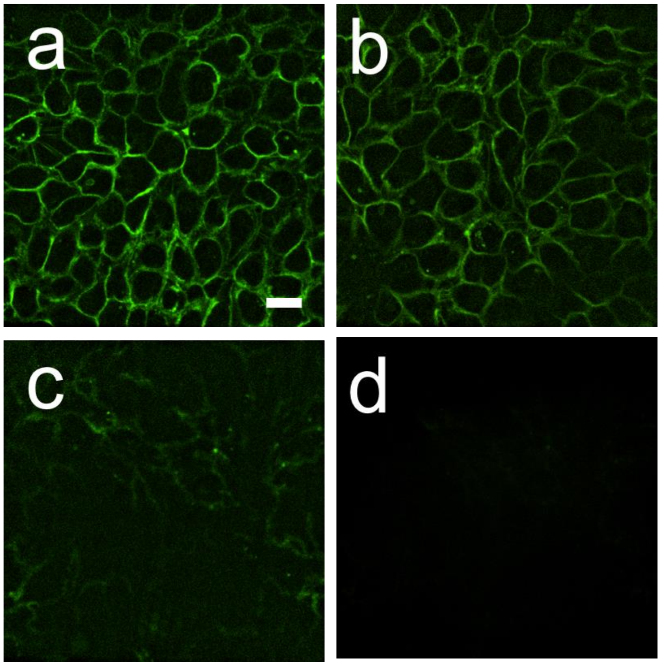

**Supplementary Figure 12. Persistence of the nqEC-DNAMeter in live cell membranes in different buffer system.**

(**a**) Representative confocal image of nqEC-DNAMeter-modified MDCK cells in HEPES-buffered saline before replacing with a complete growth medium. Scale bar, 20 µm.

(**b**) Representative confocal image of nqEC-DNAMeter-modified MDCK cells after 1.5 h incubation in HEPES-buffered saline at room temperature.

(**c, d**) Representative confocal image of nqEC-DNAMeter-modified MDCK cells after (a) 1.5 h and (b) 3 h incubation in complete growth medium at 37°C.

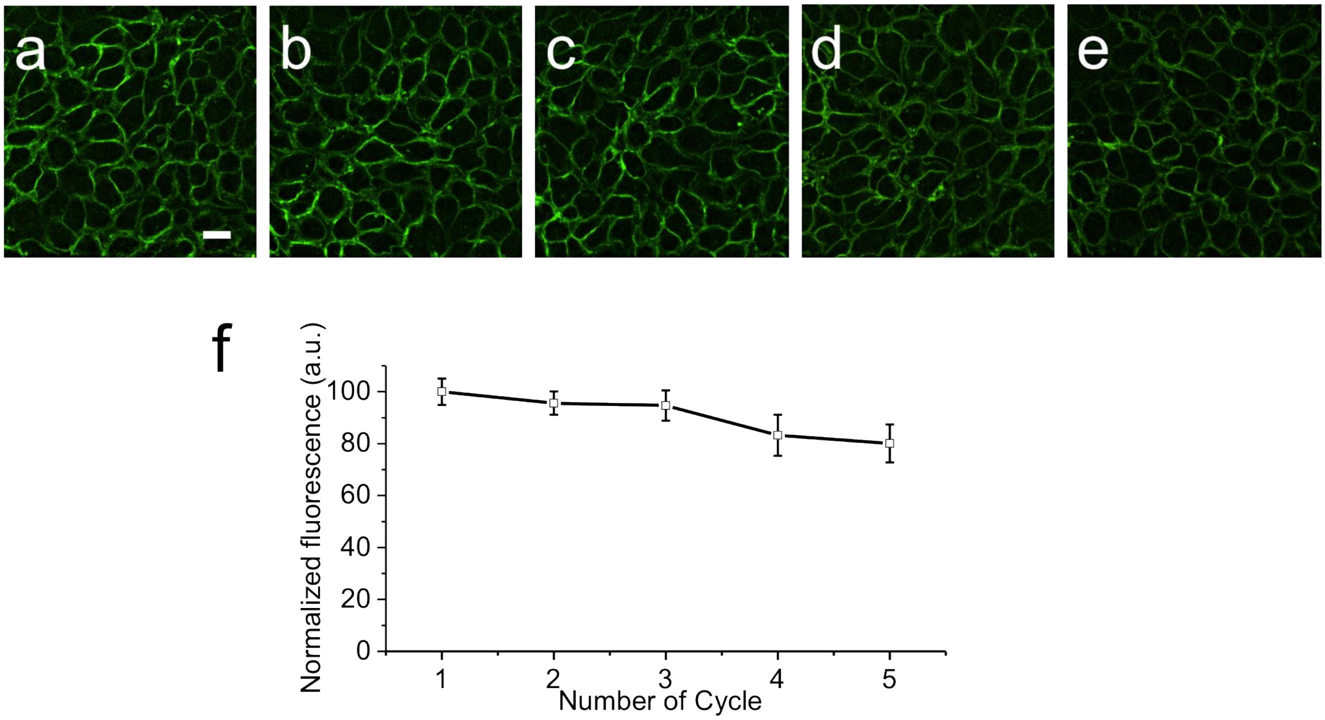

**Supplementary Figure 13. Re-insertion of the nqEC-DNAMeter into live cell membranes.**

(**a – e**) Representative confocal image of MDCK cells anchored with nqEC-DNAMeter at 0, 3, 6, 9 and 12 h in HEPES-buffered saline. After 1 h incubation with 0.2 µM nqEC-DNAMeter in HEPES-buffered saline, the MDCK cells were imaged with a 40× oil immersion objective to obtain images at 0 h (a). Then the buffer was replaced with the complete growth medium and the cells were grown at 37°C with 5% $CO_2$ for 3 h. After replacing with HEPES-buffered saline containing another 0.2 µM nqEC-DNAMeter and incubated for 1 h, the cells were imaged again to obtain images at 3 h (b). The above steps were repeated for another three times for cell imaging at 6, 9 and 12 h (c – e). Scale bar, 20 µm.

(**f**) Normalized fluorescence on the MDCK cell membranes during the above-mentioned re-insertion cycles.

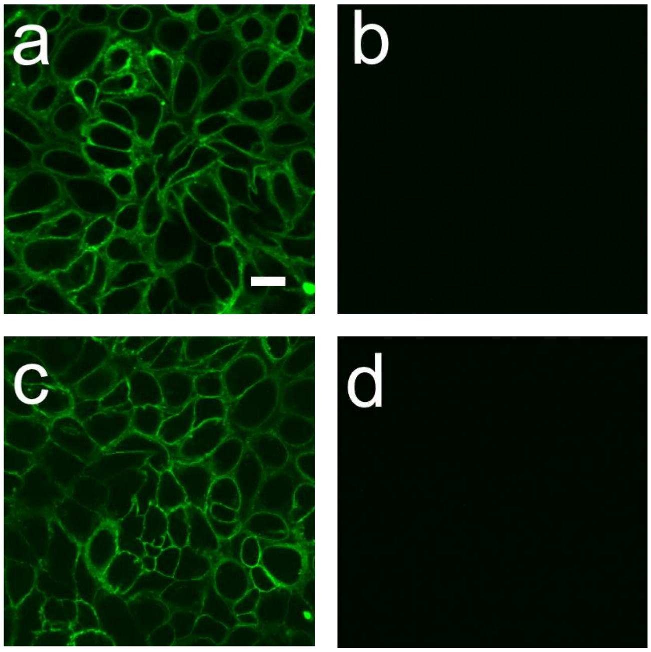

**Supplementary Figure 14. Re-insertion of the nqEC-DNAMeter into live cell membranes.**

(**a**) Representative confocal image of nqEC-DNAMeter-modified MDCK cells in HEPES-buffered saline before replacing with a complete growth medium. Scale bar, 20 µm.

(**b**) Representative confocal image of MDCK cells in HEPES-buffered saline after 12 h growth in the complete growth medium.

(**c**) Representative confocal image of cells re-inserted with 0.2 µM nqE-cad-22GCDNAMeter in HEPES-buffered saline after 12 h growth.

(**d**) Blank cell control without adding any DNA probe.

a

b

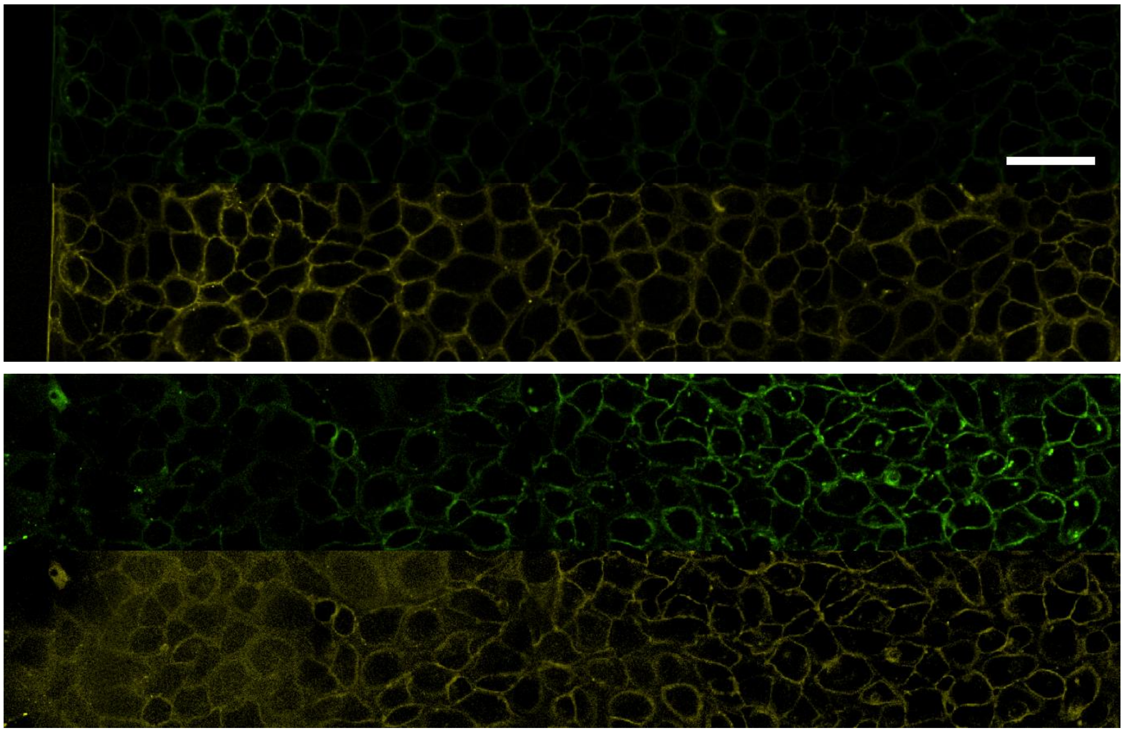

**Supplementary Figure 15. Mapping E-cadherin-mediated intercellular forces during collective cell migration.**

(**a**) Large area scan images of EC22-DNAMeter-modified MDCK monolayer cells in the green (> 4.4 pN forces) and red (reference) channels before removing the PDMS. The initial concentration of the EC22-DNAMeter was 0.2 µM. Scale bar, 50 µm.

(**b**) Large area scan images of EC22-DNAMeter-modified MDCK monolayer cells in the green (> 4.4 pN forces) and red (reference) channels after 12 h migration.

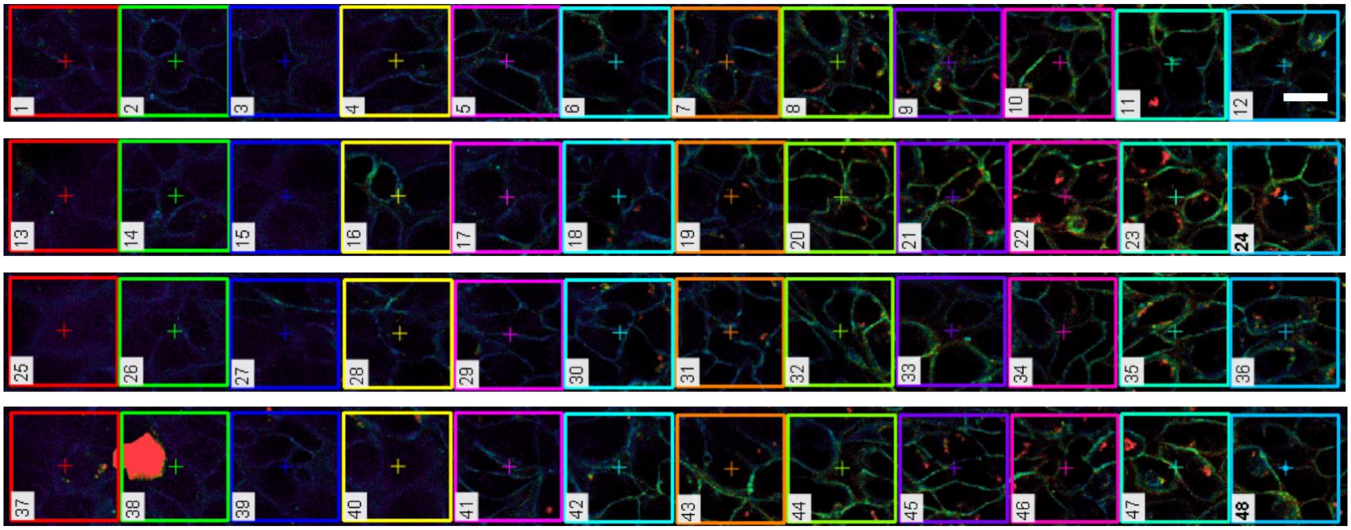

**Supplementary Figure 16. Ratiometric images for the quantitative analysis of the force distributions during collective cell migration.**

Starting from the leading edge, twelve 50 x 50 $\mu m^2$ squares were continuously selected and analyzed as a set of data. Each square was considered as a unit area. The number of pixels with positive ratios ranging from 0.75 – 3.0 was counted and plotted with their corresponding distances to the leading edge. Scale bar, 20 µm.

**Supplementary Table 1. The sequences of oligonucleotides used in this study.**

| Name | DNA Sequence (5' – 3') |
|---|---|
| Ligand strand | Cy5 –GAGTCCTCACACTTGCTTCGATTT– SH |
| 22%GC hairpin | Chol –CCCGTGAAATACCGCACAGATGCGTTTGTATAAATGTTTTTT<br>TCATTTATACTTTAAGAGCGCCACGTAGCCCAGC– QSY21 |
| 66%GC hairpin | TAMRA –TCGAAGCAAGTGTGAGGACTCTTTCTACGAGCGTTTTTT<br>TCGCTCGTAGTTTGCTGGGCTACGTGGCGCTCTT– FAM |
| Helper strand | Dabcyl –CGCATCTGTGCGGTATTTCACCCC |
| CS to 22%GC | AAAGTATAAATGAAAAAAACATTTATACAAA |
| CS to 66%GC | AAACTACGAGCGAAAAAAACGCTCGTAGAAA |
| 1HP ligand strand | HS –TTTGCTGGGCTACGTGGCGCTCTT– FAM |
| 1HP 22%GC hairpin | Chol –CCCGTGAAATACCGCACAGATGCGTTTGTATAAATGTTTTTT<br>TCATTTATACTTTAAGAGCGCCACGTAGCCCAGC– TAMRA |

The underlined sequences are expected to fold into hairpin structures.

**Supplementary Table 2. $F_{1/2}$ calculation for the 22%GC and 66%GC DNA hairpins.**

| Name | Sequence (5' – 3') | Length (mer) | $\Delta G_{fold}$ (kJ/mol) | $\Delta G_{Stretch}$ (kJ/mol) | $\Delta x$ (nm) | $F_{1/2}$ (pN) |
|---|---|---|---|---|---|---|
| 66%GC hairpin | CTACGAGCGTTTTTTTCGCTCGTAG | 25 | 32.51 | 9.5 | 8.6 | 8.1 |
| 22%GC hairpin | GTATAAATGTTTTTTTCATTTATAC | 25 | 13.05 | 9.5 | 8.6 | 4.4 |

**Supplementary Table 3. The percentage of pixels experiencing tensions at different cell–cell junctions as used in Figure 3d.**

| Number | G- / R- (<4.4 pN) | G+ / R- (4.4 – 8.1 pN) | G+ / R+ (>8.1 pN) |
|---|---|---|---|
| 1 | 50.12 | 30.92 | 18.97 |
| 2 | 52.60 | 28.04 | 19.36 |
| 3 | 63.08 | 29.18 | 7.74 |
| 4 | 46.57 | 43.87 | 9.56 |
| 5 | 76.35 | 11.56 | 12.09 |
| 6 | 72.85 | 10.78 | 16.37 |
| 7 | 55.79 | 31.25 | 12.96 |
| 8 | 39.48 | 34.78 | 25.74 |
| 9 | 45.97 | 42.72 | 11.31 |
| 10 | 45.04 | 37.98 | 16.98 |
| 11 | 77.75 | 9.76 | 12.48 |
| 12 | 60.87 | 25.76 | 13.37 |
| 13 | 63.35 | 25.71 | 10.95 |
| 14 | 68.94 | 15.02 | 16.04 |
| 15 | 52.18 | 29.10 | 18.72 |
| 16 | 53.60 | 30.74 | 15.66 |
| 17 | 65.95 | 23.32 | 10.73 |
| 18 | 37.57 | 48.74 | 13.68 |
| 19 | 73.96 | 10.15 | 15.89 |
| 20 | 67.07 | 22.57 | 10.36 |
| **Mean** | 58.45 | 27.10 | 14.45 |
| **SEM** | 12.31 | 11.48 | 4.24 |

**Supplementary Table 4. The percentage of unfolded probes experiencing tensions at different cell–cell junctions as used in Figure 3f.**

| **Number** | **G- / R- (<4.4 pN)** | **G+ / R- (4.4 – 8.1 pN)** | **G+ / R+ (>8.1 pN)** |
|---|---|---|---|
| 1 | 80.4 | 13.2 | 6.4 |
| 2 | 85.8 | 11.1 | 3.1 |
| 3 | 78.2 | 17.5 | 4.3 |
| 4 | 74.1 | 16.3 | 9.6 |
| 5 | 82.1 | 15.5 | 2.4 |
| 6 | 83.6 | 8.2 | 8.2 |
| 7 | 80.1 | 11.7 | 8.2 |
| 8 | 78.9 | 13.5 | 7.6 |
| 9 | 85.3 | 10.6 | 4.1 |
| 10 | 75.5 | 16.9 | 7.6 |
| **Mean** | 80.4 | 13.5 | 6.2 |
| **SEM** | 3.9 | 3.1 | 2.5 |

### References

[1] Abraham, M. J.; Murtola, T.; Schulz, R.; P´all, S.; Smith, J. C.; Hess, B.; Lindahl, E. *SoftwareX* **2015**, *1*, 19-25.

[2] Huang, J.; Rauscher, S.; Nawrocki, G.; Ran, T.; Feig, M.; de Groot, B. L.; Grubmu¨ller, H.; MacKerell Jr, A. D. *Nat. Methods* **2017**, *14*, 71-73.

[3] Jorgensen, W. L.; Chandrasekhar, J.; Madura, J. D.; Impey, R. W.; Klein, M. L. *J. Chem. Phys.* **1983**, *79*, 926-935.

[4] Jo, S.; Kim, T.; Iyer, V. G.; Im, W. *J. Comput. Chem.* **2008**, *29*, 1859-1865.

[5] http://www.scfbio-iitd.res.in/software/drugdesign/bdna.jsp#.

[6] Berendsen, H. J. C.; Postma, J. P. M.; DiNola, A.; Haak, J. R. *J. Chem. Phys.* **1984**, *81*, 3684-3690.

[7] Bussi, G.; Donadio, D.; Parrinello, M. *J. Chem. Phys.* **2007**, *126*, 014101.

[8] Hess, B.; Bekker, H.; Berendsen, H. J. C.; Fraaije, J. G. E. M. *J. Comput. Chem.* **1998**, *18*, 1463-1472.

[9] Darden, T.; York, D.; Pedersen, L. *J. Chem. Phys.* **1993**, *98*, 10089-10092.
